# AR coactivators, CBP/p300, are critical mediators of DNA repair in prostate cancer

**DOI:** 10.1101/2024.05.07.592966

**Authors:** Sumaira Sardar, Christopher M. McNair, Lakshmi Ravindranath, Saswati N. Chand, Wei Yuan, Denisa Bogdan, Jon Welti, Adam Sharp, Natalie K. Ryan, Matthew J. Schiewer, Elise G. DeArment, Thomas Janas, Xiaofeng A. Su, Lisa M. Butler, Johann S. de Bono, Kris Frese, Nigel Brooks, Neil Pegg, Karen E. Knudsen, Ayesha A. Shafi

**Author notes:** Corresponding Author: Ayesha A. Shafi PhD, Center for Prostate Disease Research (CPDR) 6720A Rockledge Drive, Suite 300, Bethesda, MD, 20817. Equal contribution (Sumaira Sardar, Christopher M. McNair). Disclaimer: The contents of this publication are the sole responsibility of the author(s) and do not necessarily reflect the views, opinions, or policies of Uniformed Services University of the Health Sciences (USUHS), The Henry M. Jackson Foundation for the Advancement of Military Medicine, Inc., the Department of Defense (DoD), the Departments of the Army, Navy, or Air Force. Mention of trade names, commercial products, or organizations does not imply endorsement by the U.S. Government.

## Abstract

Castration resistant prostate cancer (CRPC) remains an incurable disease stage with ineffective treatments options. Here, the androgen receptor (AR) coactivators CBP/p300, which are histone acetyltransferases, were identified as critical mediators of DNA damage repair (DDR) to potentially enhance therapeutic targeting of CRPC. Key findings demonstrate that CBP/p300 expression increases with disease progression and selects for poor prognosis in metastatic disease. CBP/p300 bromodomain inhibition enhances response to standard of care therapeutics. Functional studies, CBP/p300 cistrome mapping, and transcriptome in CRPC revealed that CBP/p300 regulates DDR. Further mechanistic investigation showed that CBP/p300 attenuation via therapeutic targeting and genomic knockdown decreases homologous recombination (HR) factors *in vitro*, *in vivo*, and in human prostate cancer (PCa) tumors *ex vivo*. Similarly, CBP/p300 expression in human prostate tissue correlates with HR factors. Lastly, targeting CBP/p300 impacts HR-mediate repair and patient outcome. Collectively, these studies identify CBP/p300 as drivers of PCa tumorigenesis and lay the groundwork to optimize therapeutic strategies for advanced PCa via CBP/p300 inhibition, potentially in combination with AR-directed and DDR therapies.

## Introduction

Prostate cancer (PCa) remains the second most common cause of male cancer mortality in the USA^1,2^. The androgen receptor (AR), a hormone-activated transcription factor, plays vital roles in the development and progression of PCa. Thus, androgen-deprivation therapy (ADT) is a standard-of-care first-line therapy for metastatic PCa. Resistance to ADT leads to almost uniformly lethal disease, termed castration-resistant prostate cancer (CRPC)^3–7^. Since the introduction of hormonal therapy ∼60 years ago^8^, progress in developing definitive treatments for aggressive disease stage has been difficult despite major advances in understanding prostate carcinogenesis and disease biology^9,10^. Recent development of abiraterone, enzalutamide, apalutamide, darolutamide, radium-223, PSMA-Lu, and cabazitaxel have improved outcome, but metastatic PCa remains a uniformly fatal disease^11–14^. Moreover, while molecular subtyping affords therapeutic benefit and improved patient survival in other tumor types, some work has been done in PCa including PAM50 subtyping and presence DNA repair alterations, but additional work remains to be achieved for more detailed subtyping in PCa^15–19^. The majority of patients with metastatic PCa are treated identically, without selection of appropriate therapeutic regimens based on tumor profile, and there is no durable therapy for metastatic disease.

Whereas local disease can be effectively treated through radical prostatectomy or radiotherapy^3,20^, non-organ confined disease presents a significant clinical challenge. First-line treatment for disseminated disease consists of androgen deprivation therapy (ADT), as PCa cells are exquisitely dependent on androgen receptor (AR) signaling for growth and survival^3,12,13,21^. This regimen is often complemented with the use of direct AR antagonists^9,10,22–26^. ADT is initially effective in most patients, and successful ablation of AR activity is validated by loss of detectable prostate specific antigen (PSA). Notably, PSA is a well-defined AR target gene, and since the protein product is secreted into the sera, it is a convenient biochemical readout of prostate-specific AR function. ADT results in a heterogeneous cellular response of tumor cell quiescence and cell death, resulting in cancer remission^9,12,23,24,27,28^. Unfortunately, this effect is transient and lasts only 2-3 years, at which time the incurable form of the disease, castration resistant prostate cancer (CRPC), emerges^3,13,14^. It is well established that this transition is largely driven by inappropriate AR reactivation despite the continuation of ADT, leading to patient morbidity^3,13,28–31^. Thus, it is critical to define alternative, complementary strategies that can act in concert with AR-directed therapeutics to suppress CRPC growth and progression.

CBP/p300 are paralogous, highly conserved histone acetyltransferases (HAT) that serve as transcriptional co-activators^32–34^. Each harbors domains that interact with sequence-specific transcription factors (including AR and c-Myc). Notably, high p300 expression has been associated with locally advanced disease and castration-resistant AR function^32–34^. Previous studies have demonstrated that CBP expression is also elevated in clinical specimens as a function of disease progression^35^. Furthermore, studies using reporter assays nominated CBP/p300 as coregulators in support of AR and c-Myc^32,33,36–40^. In PCa models, CBP/p300 expression is induced in response to androgen ablation, suggesting that CBP/p300 may support disease progression by amplifying basal AR activity in the castration setting, and thereby enhance tumor progression. Given the potential of CBP/p300 as therapeutic targets, especially for malignancies that are driven by CBP/p300-dependent transcription factors, CBP/p300 functional activities have been nominated as a possible node of intervention. Previous attempts focused on suppression of the HAT activity of CBP/p300^34,40–50^, but these strategies proved ineffective in the preclinical setting.

Here, this collaborative study shows that CBP/p300 expression is enhanced in advanced disease and associated with poor outcome. Furthermore, CBP/p300 correlate closely with AR gene expression and AR activity score in primary PCa and CRPC. By employing clinically relevant PCa models, the clinical significance of CBP/p300 expression in PCa patients as well as mechanistic evaluation of CBP/P300 transcriptional reprogramming and DNA damage response pathways were investigated. Findings revealed that CBP/p300 bromodomain suppression sensitizes to AR-dependent DNA-repair. Transcriptional mapping identified CBP/p300 as regulators of cell proliferation and DNA repair processes, which were functionally confirmed across PCa model systems. To assess relevance, exogenous challenge with genotoxic stress (utilizing *in vitro* systems, *in vivo* models, and human tumors *ex vivo*) revealed that the CBP/p300 bromodomain is required for AR-mediated DNA repair, and CBP/p300 expression is linked to DNA repair capacity in the clinical setting. Molecular analyses revealed that CBP/p300 facilitate double-strand break (DSB) repair efficiency via homologous recombination (HR) mediated DNA damage response (DDR). Congruently, CBP/p300 strongly correlated with HR gene expression in PCa patient tissue. These collective findings reveal that CBP/p300 govern rapid repair of DNA DSBs by regulating HR gene expression, thus modulating genome integrity, and promoting CRPC growth. In sum, these studies identify CBP/p300 as a driver of PCa tumorigenesis through coordinated control of critical transcriptional events and lay the groundwork to optimize therapeutic strategies for advanced PCa via CBP/p300 inhibition, potentially in combination with AR-directed therapies and DDR agents to enhance patient outcome.

## Results

### CBP/p300 expression is enhanced in advanced disease and selects for worse outcome in metastatic disease

Several PCa studies have highlighted the importance of AR-mediated DNA repair factor regulation, yet this critical facet of AR signaling is incompletely defined. In response to androgen stimulation, AR regulates a vast transcriptional network as illustrated by cistrome and transcriptome mapping. Furthermore, AR-dependent DNA repair factor regulation is a major effector of the response to DNA damage^51–54^. Thus, the AR signaling axis is a key component to further target for potential mechanistic intervention to enhance patient outcome. Importantly, CBP/p300 are paralogous, highly conserved histone acetyltransferases that serve as transcriptional co-activators for AR^55,56^. To determine the expression level and importance of CBP/p300 and AR in disease progression, hormone sensitive prostate cancer (HSPC) and castration resistant prostate cancer (CRPC) patient tissue from publicly available datasets on cBioPortal (TCGA, MSK/DFCI, (Nature Genetics 2018), and SU2C/PCF Dream Team (PNAS 2015, 2019)) were examined. In the HSPC cohort, the total alterations of patients harboring AR, CBP, and p300 alterations included 9 patients with AR alterations, 12 patients with CBP alterations, and 7 patients with p300 alterations. Additionally, in the CRPC cohort, the total alterations of patients harboring AR, CBP, and p300 alterations included 252 patients with AR alterations, 36 patients with CBP alterations, and 8 patients with p300 alterations. *AR* is amplified in both HSPC (55.6%) and CRPC (78.4%) (Fig. 1A, Supp Fig. 1A). *CBP* is characterized by 58.3% mutation and 41.7% amplification in HSPC. While *p300* is characterized by 85.7% mutation, and 14.3% amplification in HSPC (Fig. 1A). Importantly, with disease progression, patients with CRPC tumors harboring AR, CBP, and p300 alterations are characterized by frequent amplification of CBP and p300 at 83.3% and 37.5%, respectively. Similar increases in amplification of CBP/p300 are observed in additional data sets from MSK/DFCI, Nature Genetics 2018 and SU2C/PCF Dream Team Cell 2015 on cBioPortal (Supp Fig. 1B). Furthermore, CBP and p300 mRNA expression are significantly correlated (spearman = 0.67) in metastatic disease in the SU2C/PCF Dream Team cBioPortal cohort (Fig. 1B, Supp Fig 1.C) indicating that the expression of these two key AR modulators is linked in advanced disease. Additionally, CBP expression is also elevated in clinical specimens as a function of disease progression as shown in our previous study^35^. The study showed that the immunohistochemistry (IHC) analysis of CBP in HSPC needle biopsy displays lower levels of CBP compared to CRPC bone marrow transplant which exhibits a significant (p<0.001) increase in CBP expression. Subsequently, heightened levels of CBP/p300 expression are associated with unfavorable patient outcomes. Notably, survival probability significantly decreases with higher expression of CBP (p=0.05) and p300 (p=0.01) (Fig. 1C). This observed trend is further supported by cBioPortal data from the Prostate Adenocarcinoma (MSKCC, Cancer Cell, 2010) dataset (Fig. 1D, Supp Fig. 1D). Analyzing the correlation between altered AR, CBP, and p300 and disease-free survival, patients with altered AR (n=20), CBP (n=29), and p300 (n=8) exhibit significantly decreased survival probability, indicated by p-values of 9.29^-8^, 4.66e^-2^, and 1.37e^-2^, respectively. In terms of disease-free survival, patients with alterations in AR, CBP, and p300 experienced a maximum of 110, 90, and 65 months, while some patients without these alterations surpassed >160 months of disease-free survival (Fig. 1D). Importantly, the correlation of CBP/p300 with disease progression and poor patient outcome were consistently replicated in various datasets, (Supp Fig.1D). Combined, these studies implicate amplification and thus functional induction of CBP/p300 as an effector of disease progression.

**Figure 1.**
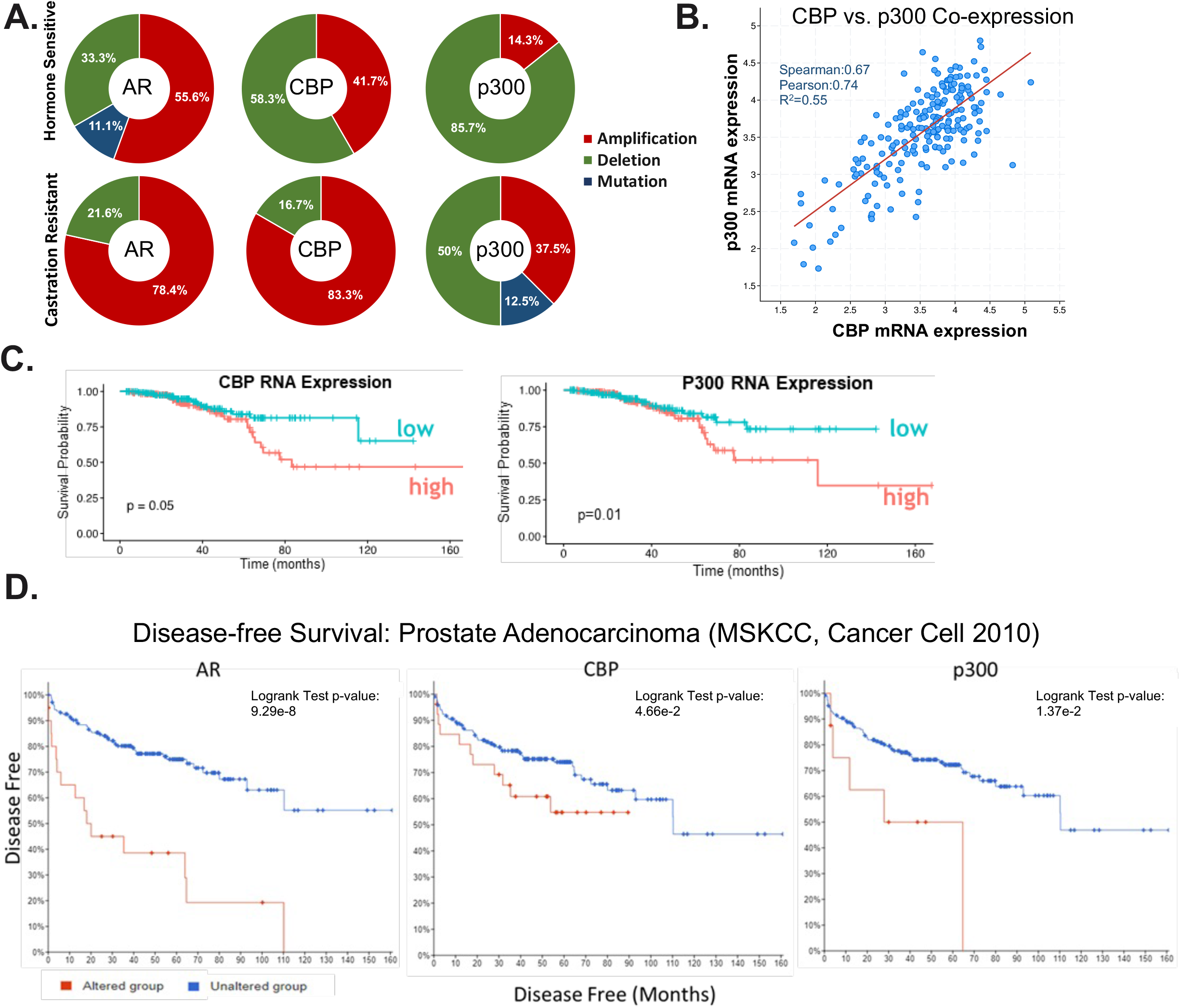
Altered CBP/p300 expression correlates with aggressive disease. **A.** Frequency of AR, CBP, and p300 gene alterations (i.e. amplifications, mutations, and/or deletions) in primary and metastatic PCa datasets from cBioPortal – TCGA Firehose Legacy and SU2C/PCF Dream Team (PNAS 2019) datasets. Hormone sensitive PCa total alterations AR n=9, CBP n=12, and p300 n=7, and castration-resistant PCa AR n=252, CBP n=36, and p300 n=8. **B.** Correlation of CBP and p300 mRNA expression in metastatic PCa patient tumors from SU2C/PCF Dream Team (PNAS 2019) data set from cBioportal. **C**. CBP and p300 are highly expressed in primary PCa (top 10% expressed genes) and mCRPC (top 25% expressed genes) and are associated with poor outcome. **D.** cBioportal analyses for disease-free survival from prostate adenocarcinoma data set (SKCC, Cancer Cell 2010).

### CBP/p300 bromodomain suppression enhances response to standard of care therapeutics

Given the potential of CBP/p300 as therapeutic targets, especially for malignancies driven by CBP/p300-dependent transcription factors, CBP/p300 functional activities have been proposed as a conceivable target of intervention. Previous attempts focused on suppression of the HAT activity of CBP/p300^34,40–50^, but these strategies proved ineffective in the preclinical setting. Our initial study^35^ utilizing CCS1477 (Inobrodib) that selectively targets the CBP/p300 bromodomain, which recognizes acetylated lysine residues, demonstrated CBP/p300 bromodomain inhibition impacts AR and c-Myc signaling, resulting in decreased tumor growth in CRPC^35,36,42,45^. As previously reported, CCS1477 binds to the bromodomain of CBP/p300 and inhibits tumor growth by disrupting AR signaling *in vitro*, *in vivo*, and *ex vivo*^35^. In the present study, we aimed to further elucidate the mechanism(s) of action of CBP/p300 in PCa and explore the potential clinical application of CCS1477. First, we validated the effect of CBP/p300 bromodomain inhibition by exposing CRPC cell model systems to 1 μM CCS1477 over time. CRPC cells exposed to CCS1477 were harvested at 0, 4, 24, and 48 hours to determine the impact on the AR signaling axis via AR, AR-SV (splice variants – AR-V7), c-MYC, CBP, and p300 expression. Consistent with our previous study, AR isoforms and c-MYC protein expression decreased over time while CBP and p300 expression remained stable (Fig. 2A). This validated previous findings that CBP/p300 inhibition decreased AR signaling suggesting altered function of CBP/p300 without impacting protein expression. The biological effect of CBP/p300 on cell cycle progression in CRPC was assessed in multiple cell models, wherein CBP/p300 bromodomain inhibition resulted in decreased S phase and G2/M arrest over time (Fig. 2B, Supp Fig. 2A). As an important cellular checkpoint, G2/M arrest is associated with DNA damage repair (DDR). In accordance, key DDR and cell cycle markers including *c-MYC*, *TP53*, and *CDKN1A* mRNA expression decreased with CCS1477 at 0, 12, 24, and 48 hours in CRPC models (Supp Fig. 2B). Thus, tumor-associated AR coactivators, CBP/p300, appear to be essential for cellular proliferation. To examine the biological relevance, combinatorial studies with standard-of-care (SOC) therapies and CBP/p300 inhibition were performed for downstream cell survival assessment. Specifically, SOC therapies such as irradiation (IR), cisplatin (platinum-based therapy), olaparib (PARP inhibitor), and doxorubicin were assessed in combination with increasing concentrations of CCS1477. Importantly, combination therapy at the highest doses were the most lethal in CRPC models (Figs. 2C-E, Supp Figs. 2C-E). Specifically, the lowest relative cell growth is observed in CRPC cells treated with CCS1477 and higher doses of SOC with enhanced combination index achieved in these combinatorial studies indicating synergy (Figs. 2C-D and Supp Figs. 2C-D). The IC50 values decreased 5-10-fold from single agent (CCS1477) to combination with SOC in 22Rv1 and C4-2 cells (Fig. 2D). To understand the effect of the combination therapy on the AR signaling axis, AR, c-MYC, CBP, and p300 protein expression was examined in CRPC cell lines treated with 5Gy IR, 1μM cisplatin, 2.5 μM olaparib, and 0.5 μM doxorubicin alone and in combination with CCS1477 (Fig. 2E, Supp Fig. 2E). Consistently, in both cell lines, AR and c-Myc expression decreased further in combination treatment than with single agent. Interestingly, CBP/p300 expression varies between the different SOC therapies alone compared to combination. Notably, p300 protein expression decreases with CCS1477 + 5Gy IR in 22Rv1 and C4-2 cells compared to 5Gy IR alone. In sum, CCS1477 demonstrates promising potential as a novel small molecule inhibitor of CBP/p300 through a synergistic effect in combination with existing therapies for CRPC.

**Figure 2.**
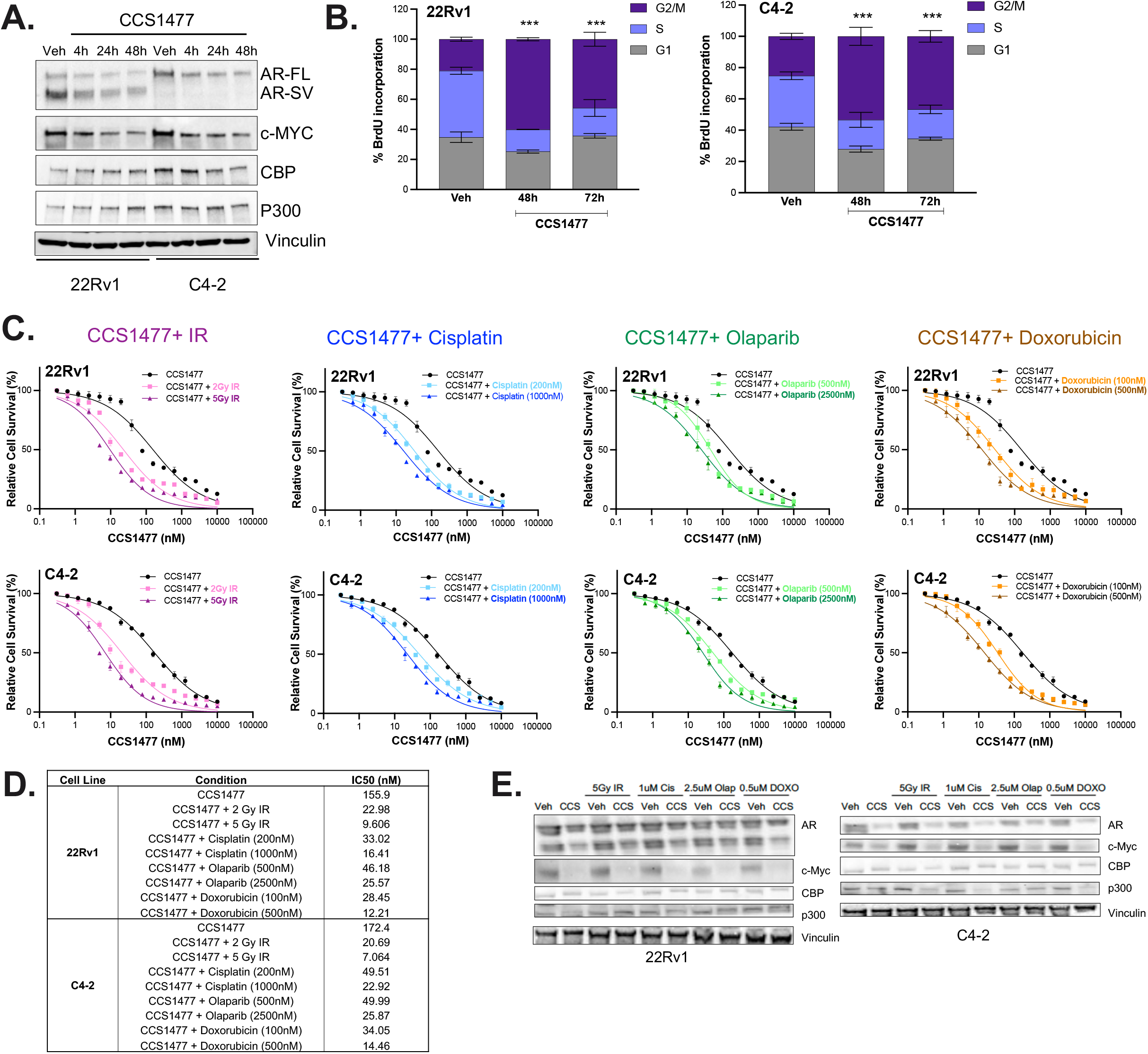
CCS1477 sensitizes CRPC cells to AR-dependent DNA repair. **A-B.** 22Rv1 and C4-2 cells treated with CBP/p300 inhibitor (CCS1477 – 1uM) for increasing time (4-72 hours). **A.** Cells were harvested and analyzed for protein expression of AR, AR-V7, c-MYC, CBP, p300, and Vinculin. **B.** Cells were harvested and analyzed for cell cycle analyses with flow cytometry. **C-D.** 22Rv1 (top) and C4-2 (bottom) cells were treated with increasing dosages of indicated DDR agents (IR, cisplatin, olaparib, or doxorubicin) with or without CCS1477, and drug sensitivity assays were performed on Day 5 using Picogreen. **D.** Non-linear regression analyses were performed to determine the IC_50_ values. **E.** WB analyses of AR, CBP, p300, c-MYC, and Vinculin in response to IR, cisplatin, olaparib, and doxorubicin in 22Rv1 and C4-2 cells. n=3, *p<0.05, **p<0.01, & ***p<0.001.

### CBP/p300 bromodomain inhibition reduces the AR and CBP/p300 cistromic landscape

These remarkable responses strongly underscore the importance of further understanding the tumorigenic role of the CBP/p300 bromodomain in cancer progression. However, genome-wide understanding of CBP/p300 function is not yet fully defined in PCa. Thus, the mechanism through which CBP/p300 influences aggressive disease was assessed. AR function was evaluated via chromatin immunoprecipitation followed by DNA sequencing (ChIP-Seq) with CCS1477. CBP/p300 cistrome mapping in CRPC, along with AR, using a stringent cutoff identified 31137, 35775, and 3581 binding sites of AR, CBP, and p300, respectively in 22Rv1 cells. Interestingly, in the presence of CBP/p300 bromodomain inhibition, the cistromic landscape is reduced to 19300, 11920, and 1660 AR, CBP, and p300-bound sites, respectively (Fig. 3A, Supp Figs. 3A-B). This decrease in AR, CBP, and p300 binding suggests a putative loss of function in AR function when CBP/p300 is inhibited, likely contributing to the biological changes seen in Figure 2.

**Figure 3.**
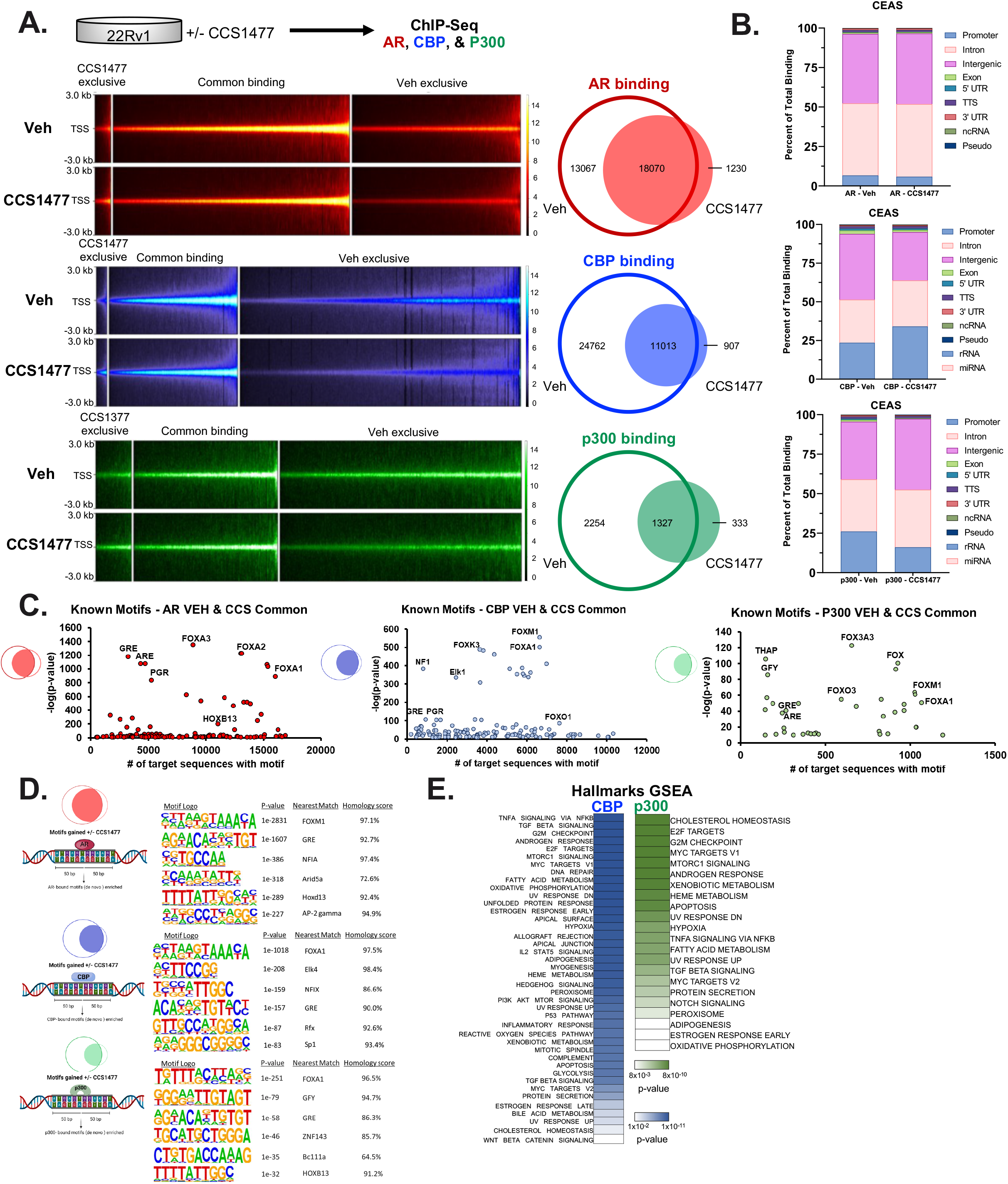
CBP/p300 inhibition drastically reduces the AR, CBP, p300 cistromic landscape. **A.** ChIP-Seq of intensity plots of 22Rv1 control and CCS1477 treated cells. AR, CBP, and p300 were immunoprecipitated in CCS1477 exclusive, common binding, and VEH exclusive in a 6-kb region. **B.** Percent of total binding of overlap of common bindings sites of AR, CBP, & p300 in CCS treated 22Rv1 cells. **C.** Known motif of AR, CBP, and p300 of common binding in 22Rv1 CCS1477 treated cells. **D.** Motifs of binding regions of AR, CBP, and p300 visualized for top 6 highest p-values. **E.** Hallmarks GSEA pathways regulated by CBP and p300 in CCS1477 treated 22Rv1 cells.

Decreased AR, CBP, and p300 have previously shown to change promotor binding^32,57,58^. As CCS1477 treatment decreased AR function and cistromic landscape, genomic annotation of AR, CBP, and p300 binding sites were performed using cis-regulatory element annotation system (CEAS) package to elucidate a description of impacted regulatory sites (Fig. 3B). As shown in Figure 3B, CBP/p300 bromodomain inhibition did not significantly alter AR regions of binding, with majority of binding observed at intronic (45%) and intergenic (44%) regions, followed by promoters (6%) with or without CBP/p300 inhibition (Fig. 3B – top). Interestingly, CCS1477 treatment shifted the regions of binding for CBP from predominately intergenic (42%), intronic (28%), and promoter (24%) to predominantly promoter (34%), intergenic (31%), and then intronic (29%) regions (Fig. 3B – middle). Conversely, inhibition of CBP/p300 bromodomain enhanced p300 binding towards intergenic (from 36% to 45% with CCS1477) and intronic (from 33% to 37% with CCS1477) regions. Additionally, CCS1477 treatment results in decreased binding to the promoter (26% to 16% with CCS1477) regions (Fig. 3B – bottom), suggesting that p300 binding shifts towards intergenic and intronic regions in CRPC following CBP/p300 inhibition. This suggests that CBP and p300 might have distinct regulatory activity which were next assessed with motif analyses to determine potential binding partners. Specifically, to discern potential transcription factors associated with AR, CBP, and p300, known-motif analysis was conducted using a broad window around the center of binding (1 KB) (Fig. 3C, Supp Fig. 3C). Expected enrichment of AR binding elements (ARE, GRE) and key pioneer factors (FOXA1, FOXA2, FOXM1) was complemented by enrichment of cancer-associated transcription factors of PCa relevance, including several with oncogenic activity (e.g., NF1, HOXB13, ETS factors)^59–64^. Specifically, enriched motifs correspond to components of the FoxA1/AR complex or are directly driven by AR to promote AR signaling, cell proliferation/invasion, metabolic rewiring, and ultimately tumor growth (Fig. 3C - right). Analysis of known CBP binding motifs showed prevalent binding to FOXA1, FOXM1, FOXO1, ELK1, and NF1 (Fig. 3C - middle). Similarly, analyses of known p300 binding motifs depicted binding to the FOXM1, FOXA1, FOXO3, FOXA3A, and THAP (Fig. 3C - left). To further investigate the potential mechanisms of CBP/p300 function, *de novo* motif analysis was assessed using a window of 50 bp adjacent to the center of binding (Fig. 3D). In addition to motifs for Forkhead Box proteins (e.g., FOXM1), multiple motifs of PCa relevance were enriched proximal to AR binding, including several factors elevated in PCa, linked with androgen-associated cancer growth, interleukin regulation, and cancer progression (e.g., NFIA, GRE, and Arid5a)^65,66^ (Fig. 3D – top). Importantly, additional motifs were enriched proximal to CBP and p300 binding, including another Forkhead Box protein (FOXA1), a key pioneer factor pivotal to PCa progression (Fig. 3D)^60,63^. Further, CBP was enriched for PCa-associated transcription factors, including ETS-factor, Elk4 and SP1^62,64,67–70^. Conversely, in addition to FOXA1, p300 was enriched for mitochondrial factor (GFY), zinc finger proteins (Bc111a, ZNF143), and PCa-associated transcription factors (HOXB13)^71–78^. Pathway analysis utilizing Hallmarks and KEGG gene set enrichment analyses (GSEA) identified that CBP and p300-bound sites impact key hallmarks of cancers including cell cycle, DNA repair, and metabolic signaling (Fig. 3E, Supp Fig. 3D-E). Transcriptional signatures also identified in this study highlight pathways of functional importance. In sum, these observed enrichments and related pathway analyses (Fig. 3E, Supp Fig. 3D-E) provide the first insight into genome-wide CBP/p300 activity in CRPC models and reveal potential exclusive oncogenic functions of the coactivator partners beyond AR signaling axis. These data provide insights into the potential mechanism driving AR signaling and CBP/p300 cofactor alterations in CRPC.

### CBP/p300 govern transcriptional networks critical for response to DNA damage

The biological significance of the AR signaling axis in driving progression to CRPC is well established^6,12,21,22,28,30,35,40,41,43,45,48,79^, and thus AR remains the most preferred target in treating PCa. However, with the observation that AR co-activators CBP/p300 are associated with poor outcomes in metastatic PCa^35^ (Fig. 1), coupled with linkage of the newly identified CBP/p300 cistrome to cancer-promoting pathways (Fig. 3), it is now cogent that CBP/p300 play pivotal and potentially targetable roles in PCa. Previous studies identified that CBP/p300 inhibition via CCS1477 resulted in significant transcriptional alterations including androgen response and MYC target pathways^35^ (Supp Fig. 4A). In this study, we performed genome-wide assessment of the CBP- and p300-sensitive transcriptional networks utilizing newly generated doxycycline-regulated isogenic paired model systems of inducible *CBP* and *p300* knockdown. As shown, ∼80% ablation of CBP and p300 protein was achieved after shRNA induction (Fig. 4A - top), subsequent to which the whole-transcriptome analysis of CBP and p300 individually was assessed in human CRPC models (Fig. 4A - bottom). Principal component analyses (PCA) indicated a high level of concordance between biological replicates as shown in sample clustering within treatment groups with strong consistency amongst biological replicates (Supp Fig. 4B). Major transcriptional changes were observed (1831 upregulated, 2785 downregulated genes) after CBP knockdown (adjusted *p*-valuelJ<lJ0.001, fold change > 2) and (3219 upregulated, 3459 downregulated genes) after p300 knockdown (adjusted *p*-valuelJ<lJ0.001, fold change > 2) indicating that CBP and p300 influence large gene networks. Gene set enrichment analysis (GSEA) (Fig. 4B, Supp Fig. 4C) revealed that CBP/p300 govern transcriptional programs of oncogenic relevance, including those involved in DNA replication, cell cycle regulation, and multiple DNA repair processes. The biological effect of CBP and p300 on cell cycle progression in CRPC was assessed in multiple distinct isogenic pairs, wherein CBP and p300 individual ablation resulted in G2/M arrest (Fig. 4C, Supp Fig 4D) in the presence of genotoxic insult, i.e., irradiation (IR). These studies reveal a requirement of CBP/p300 for cell cycle progression in CRPC. Thus, tumor-associated CBP/p300 appear to be essential for cellular proliferation thus further highlighting a therapeutic window.

**Figure 4.**
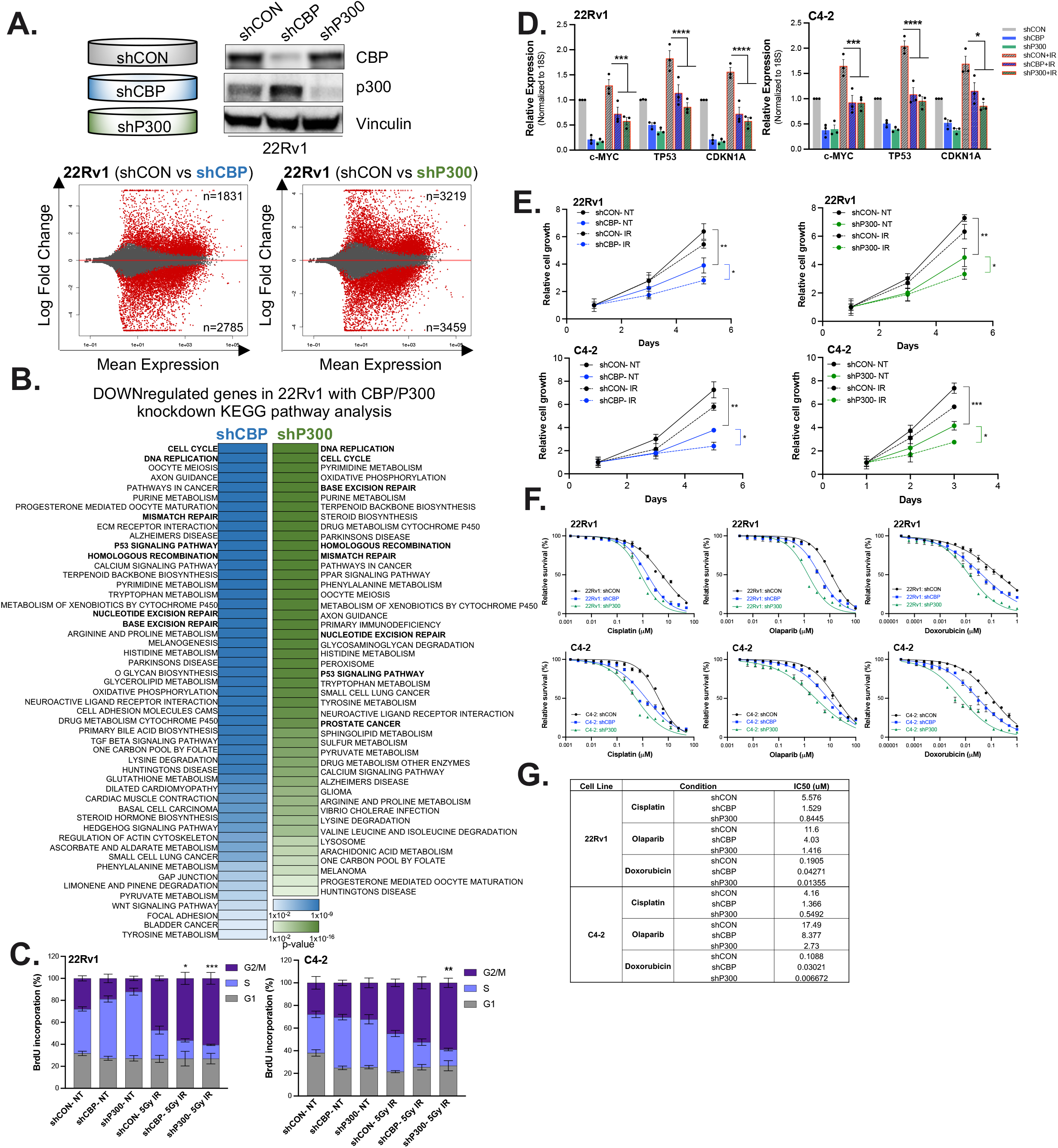
CBP/p300 acts in concert to alter the global transcriptome in PCa cells and regulate DDR. **A.** 22Rv1-shCON, shCBP, and shP300 cells were treated with doxycycline to induce knockdown, and RNA Seq analysis was performed in quadruplet samples. MA plot depicts gene expression modulation with the number of significant upregulated and downregulated transcripts with p<0.001 and fold change (FC)> 2.0. **B.** GSEA of RNA-Seq (KEGG Pathways) identified deenriched pathways with CBP and p300 knockdown treatment in 22Rv1 cells using FDR<0.25. **C-E.** 22Rv1-shCON, -shCBP, and –shp300 cells were treated with doxycycline and 5Gy IR. **C.** Flow analysis was performed to determine changes in cell cycle progression. Cells were harvested, and mRNA (**D**) was isolated. Changes in *c-MYC*, *TP53*, and *CDKN1A* (*p21*) and *18S* mRNA expression. **E.** 22Rv1-shCON, -shCBP, and –shp300 cells were treated with doxycycline to knockdown CBP and p300. Cells were also treated with IR (5Gy) at Day 1. Cells were harvested for growth assays analyses on Days 1, 3, and 5 using Picogreen. **F.** 22Rv1 (top) and C4-2 (bottom) cells were treated with increasing dosages of indicated DDR agents (Doxorubicin, Cisplatin, or Olaparib) and drug sensitivity assays were performed on Day 5 using Picogreen. **G.** Non-linear regression analyses were performed to determine the IC_50_ values. n=3, *p<0.05, **p<0.01, & ***p<0.001.

Downstream analyses demonstrated the biological relevance, as evidenced by a 1.6– 1.8-fold reduction in surviving cells after exposure to 5Gy IR in CBP-depleted and p300- depleted cells (Fig. 4E) and decreased cell growth after DNA damage from SOC (cisplatin, olaparib (PARPi), and doxorubicin) treated cells with CBP/p300 ablation (Fig. 4F-G). Specifically, the lowest relative cell growth is observed in CRPC cells treated with single agent SOC in combination with CBP and p300 knockdown (Figs. 4E-G). The IC50 values decreased 2-5-fold from single agent (SOC) to combination with CBP-depletion or p300-depletion in 22Rv1 and C4-2 cells (Fig. 4G). These findings reveal new insight into CBP/p300 function as a modulator of cell cycle checkpoint control and cell proliferation in response to DSB and implicate CBP/p300 in promoting cancer cell survival.

### CBP/p300 modulates DNA repair factor expression and homologous recombination

Since both CBP and p300 expression is elevated with disease progression and associated with worse clinical outcomes (Fig. 1), combined with molecular observations that CBP/p300 perturbations result in transcriptional rewiring influencing DNA repair (Figs. 3-4), unbiased computational strategies were employed to discern the potential mechanism of action by which CBP/p300 influence DNA damage response. To prioritize the genes for downstream analyses, genes significantly altered with stringent cutoffs (p<0.01, fold change >2.0) were compared between CBP and p300 ablation (i.e., shCBP and shP300 conditions) (Fig. 5A). This overlay identified 3000 genes in common with both CBP and p300 knockdowns in 22Rv1 cells. Next, these 3000 genes were compared to DNA damage repair (DDR) genes using MSigDB, which have been previously curated and published^80^. This resulted in a list of CBP/p300- regulated DDR genes, which were further grouped into the main DDR pathways including HR (homologous recombination), BER (base excision repair), MMR (mismatch repair), NER (nucleotide excision repair), and NHEJ (non-homologous end joining) to determine which DDR pathway is most significantly regulated by CBP/p300. These analyses revealed potentially critical roles for CBP/p300 in regulating factors associated with HR (23% of CBP/p300-regulated genes), BER (19%), MMR (15%), NER (14%), and NHEJ (1%). Concurrently, transcriptional profiling following treatment with CCS1477 revealed that CBP/p300 regulates HR factors (Supp Fig. 5B). CBP/p300 ablation and inhibition significantly impacted HR-mediated repair as functional CBP/p300 signaling is required for expression of major HR factors, including *BRCA2, POLD2, RAD51,* and *XRCC2* (Fig. 5B, Supp Fig 5B). These findings suggest that CBP/p300 modulate factors associated with HR-mediate DNA damage, impacting disease aggressiveness and response to standard-of-care.

**Figure 5.**
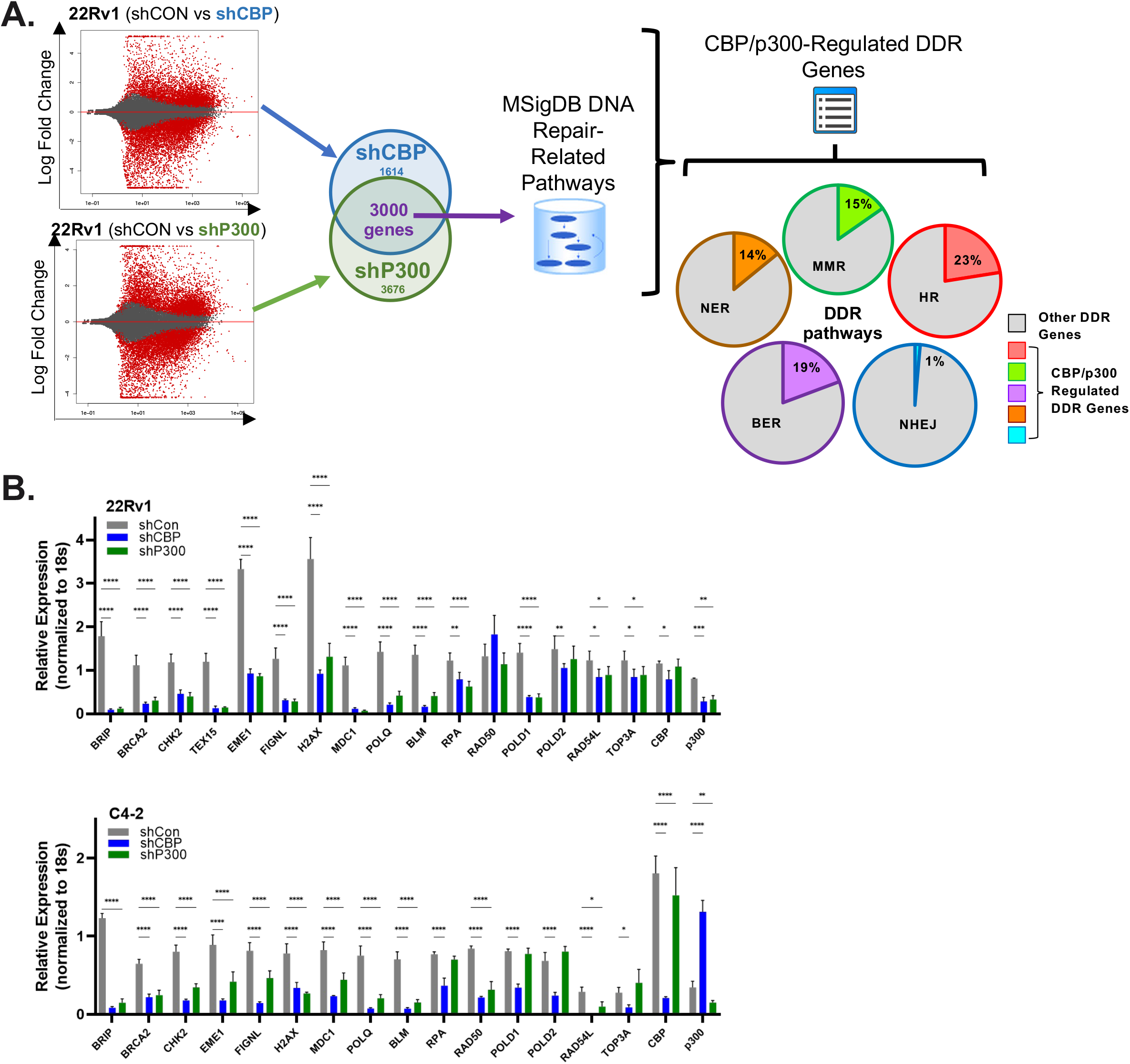
CBP/p300 inhibition impacts double-strand DNA break repair and decreases HR efficiency. **A.** Schematic describing the comparison of RNA-Seq data sets. Briefly, common transcriptions with p<0.001 and fold change (FC) > 2.0 were identified and then organized into specific DDR pathways. **B.** Validation of HR gene targets in CRPC cells with shCON, shCBP, and shp300 CRPC models (22Rv1 and C4-2).

### CBP/p300 attenuation decreases HR efficiency

An HR reporter assay was conducted by utilizing a well-established U2OS cell line to read HR repair efficiency through revived GFP expression^81,82^ (Fig. 6A – Top). CBP/p300 inhibition with 2.5 µM CCS1477 decreased HR repair by (8-fold) when compared to ATM inhibition (6-fold change) (Fig. 6A – Middle). Similar results were observed with individual and combined knockdown of CBP and p300 as compared to ATM inhibition. Overall, all inhibited cells had significantly (p<0.001) decreased HR efficiency than control cells (Fig. 6A – Bottom). These findings nominate CBP/p300 as positive effectors of HR. To assess the impact of these findings on PCa, CBP/p300 isogenic pairs with individual knockdown of CBP and p300 in the presence of genotoxic insult, IR, were assessed for impact on HR target gene expression. HR factors including, *ATM*, *CHEK2*, and *RAD50,* mRNA expression was significantly decreased in knockdown cells with genotoxic insult (Fig. 6B). To further validate these findings, 22Rv1 and C4-2 cells were treated with CCS1477 and genotoxic insult. HR factors, including *ATM*, *CHEK2*, and *RAD50* mRNA expression decreased (p<0.001) compared to IR conditions (Fig. 6C - Top). Similar results were observed in C4-2 cells (p<0.001) (Fig. 6C – Bottom and Supp Fig. 6A). Concordant changes in pATM and pCHK2 protein levels were observed as a result of CBP/p300 inhibition (Fig. 6D, Supp Fig. 6B). In sum, these findings show that CCS1477 destabilizes the HR mediated repair pathway through CBP/p300 attenuation (e.g., bromodomain inhibition and genomic knockdown).

**Figure 6.**
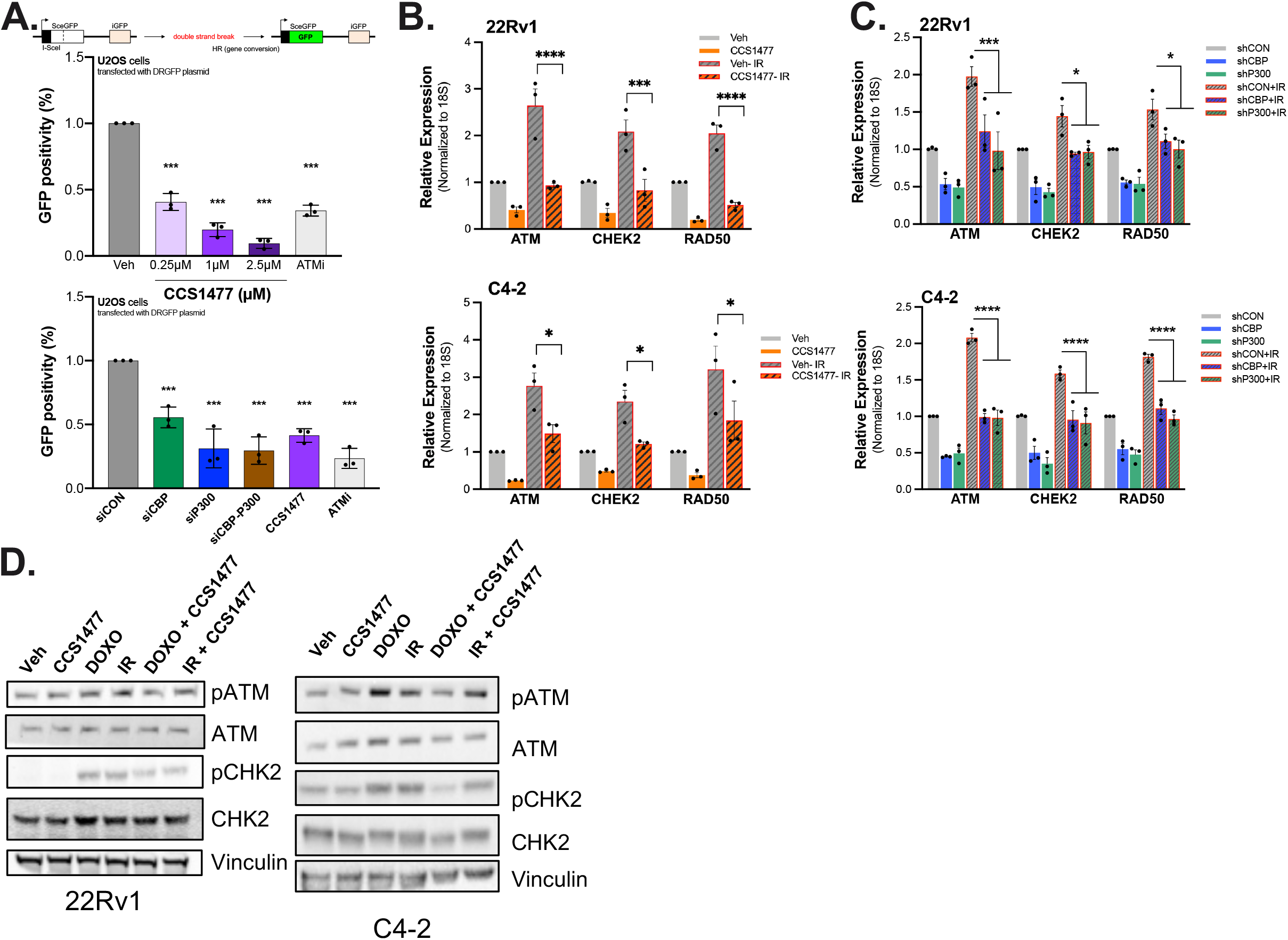
CBP/p300 impacts double-strand DNA break repair and decreases HR efficiency. **A.** U20S-DR-GFP cells were treated with increasing doses of CCS1477. CBP and p300 was knocked down in U20S-DR-GFP cells for 72 hrs via siRNA. Cells were treated with ATM inhibitor for 24 hrs. Cells were harvested for flow cytometry. **B-D.** Changes in HR factors (*ATM*, *CHEK2*, and *RAD50*) and *18S* mRNA and protein expression were analyzed with CBP/p300 attenuation (via shRNA or CCS1477 treatment) and 5 Gy IR for 24 hours. **B.** 22Rv1 & C4-2-shCON, -shCBP, and –shp300 cells were treated with doxycycline to knockdown CBP and p300. Then cells were treated with 5Gy IR. Cells were harvested and mRNA was isolated. **C.** Changes in *ATM*, *CHECK2*, *RAD50*, and *18S* mRNA expression were analyzed in CBP/p300 inhibitor treated CRPC cells. **D.** Changes in HR factors protein expression were analyzed with CBP/p300 inhibitor with or without genotoxic insult of 5 Gy IR or 10 nM Doxorubicin treatment for 24 hours. n=3, *p<0.05, **p<0.01, & ***p<0.001.

### Targeting CBP/p300 impacts HR-mediated repair and patient outcome

CBP/p300 inhibition holds therapeutic potential in combination with first-line therapy drugs (Figure 2). Therefore, the impact on HR-mediated repair was assessed in patient samples. Analysis of key HR genes with CBP/p300 attenuation (inhibition or knockdown) in CRPC cell lines were compared, and top altered HR genes were identified for impact in PCa patients (Fig. 7A). Importantly, CBP/p300 inhibition in both CRPC models (22Rv1 and C4-2) and isogenic pairs with knockdown revealed that CBP/p300 regulate key HR factors. Next, to further examine the translational relevance, HR genes (*ATM, MRE11A*, and *RAD50*) were compared for co-occurrence with *CBP* and *p300* in human data set from cBioportal (SU2C/PCF Dream Team (PNAS 2019) (Fig. 7B). Correlations for all three HR genes were statistically significant (p <0.001), highlighting HR pathway as a strong mode of DDR in patients with elevated CBP/p300. Furthermore, xenograft models of 22Rv1 cells undergoing 28-days treatment with 20 mg/kg CCS1477 displayed a significant decrease in HR target genes (*MRE11A, RAD50*, *CHEK2*, and *BRCA2*) (Fig. 7C, Supp Fig. 7A) supporting *in vitro* findings. To assess the potential for therapeutic resistance, impact of long-term treatment with CBP/p300 inhibition was evaluated using patient derived xenografts (PDX – CP50c, previously published^35^) over a period of 8-days and when tumors reached 300% of their original growth (long-term) of CCS1477 (Fig. 7D). Importantly, short-term CCS1477 treatment decreased HR-mediated repair as measured via expression levels of HR genes (*MRE11A* and *RAD50*). Moreover, CBP/p300 expression co-occurred with additional HR factors (*ATM, RAD51,* and *RAD54L*) (Supp Fig. 7B). Intriguingly, long-term CBP/p300 inhibitor treatment caused an increase in HR genes, displaying potential resistance to CCS1477 over time (Fig. 7D). Finally, to gain a better understanding of CCS1477 treatment on clinical tumor growth, patient derived explants (PDEs) were utilized^83^. Briefly, in this *ex vivo* system, tumor and adjacent non-neoplastic samples from primary PCa patients undergoing radical prostatectomy were placed on dental sponges and treated with increasing concentrations of CCS1477 (1-5 µM) (Fig. 7E-F). Increasing doses of CCS1477 significantly decreased (p<0.05) tumor growth as measured by Ki67 staining, a marker of proliferation (Fig. 7E). Furthermore, PDEs were analyzed for expression of HR genes with CBP/p300 inhibition (Fig. 7F). Consistent with the *in vitro* cell line and *in vivo* xenograft and PDX studies, decreased HR gene expression is observed in *ex vivo* PDEs with CBP/p300 bromodomain inhibition. Importantly, these findings support the interplay of the CBP/p300 axis and HR-mediated repair. Analysis of patients treated with CCS1477 indicate that CBP and p300 co-occur with HR targets, *ATM*, *RAD50*, *RAD51*, and *RAD54L*, in PCa patients (Supp Fig. 7B). Combined, these findings identify CBP/p300 attenuation as drivers of HR-mediated anti-tumorigenic signaling in PCa and identify a potential novel target to enhance patient outcome (Fig. 7G).

**Figure 7.**
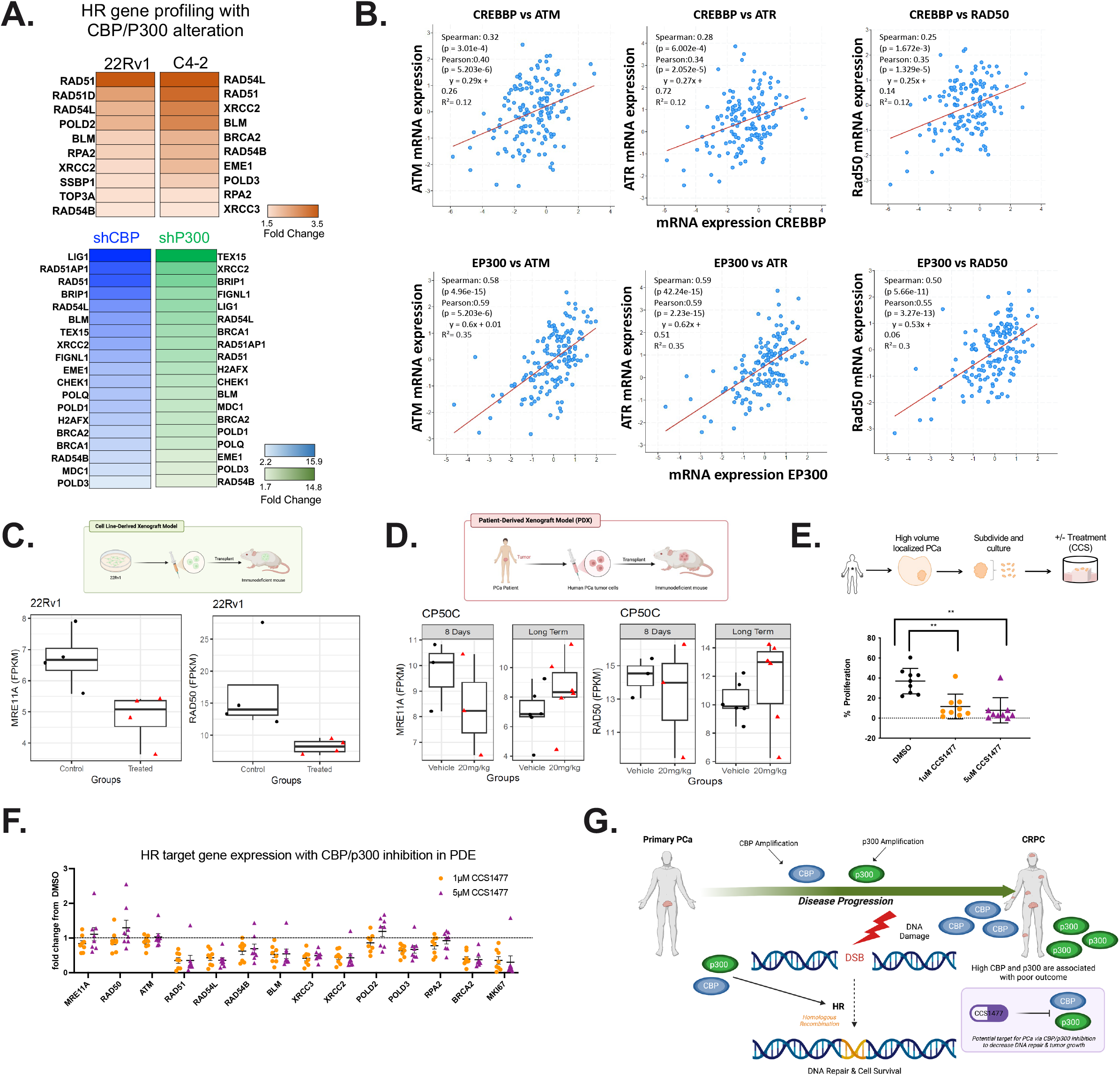
CBP/p300 inhibition mediates HR repair and impacts patient outcome. **A.** HR gene profiling in 22Rv1 and C4-2 cells with CBP/p300 inhibition (CCS1477) or CBP/p300 knockdown in inducible cell models. **B.** Correlation of HR genes (ATM, ATR, and RAD50 with CBP and p300 mRNA expression in patient cohorts from SU2C/PCF Dream Team (PNAS 2019) data set from cBioportal. **C.** HR expression in 22Rv1 xenografts with CCS1477 (20mg/ml) and harvested to examine HR gene expression. **D.** Patient-Derived Xenografts (PDX) prostate cancer tumor tissue treated with CCS1477 (2-mg/mL) for 8 weeks and then longer. Harvested, RNA isolated, and analyzed for HR gene expression. **E.** Patient-Derived Explants (PDE) prostate cancer tumor tissue treated with CCS1477 (1 uM and 5 uM) for 48 hours and analyzed for Ki67 positivity. **F.** HR expression in PDEs with CCS. **G.** Module summarizing findings. n=3, *p<0.05, **p<0.01, & ***p<0.001. n>3, *p<0.05, **p<0.01, & ***p<0.001.

## Discussion

CRPC remains an incurable disease stage with ineffective treatments options to avoid lethality. In this study, we elucidate the mechanism of CBP/p300 inhibition functioning through HR, to potentially enhance therapeutic targeting of CRPC. We previously established CBP and p300 attenuation via bromodomain inhibition and isogenic knockdown models regulate AR signaling in a c-MYC manner in CRPC *in vitro* and *in vivo* models^35^. Now, this study further illuminated the mechanism of action of CBP/p300 inhibition which resulted in decreased PCa growth by directly impacting HR-mediated DNA repair. Key findings demonstrate that: (i) CBP/p300 expression increases as disease progresses to CRPC and is associated with poor patient outcome; ii) Inhibition of the CBP/p300 bromodomain enhances response to SOC therapies; iii) Cistromic landscape of CBP and p300 is drastically reduced with CBP/p300 inhibition; iv) CBP/p300 governs pathways critical for the response to DNA damage via transcriptional regulation; v) CBP/p300 regulates HR-mediated repair *in vitro*, *in vivo*, and *ex vivo*; vi) Targeting CBP/p300 impacts HR-mediated repair; and vii) CBP/p300 are strongly associated with HR factor expression and poor outcome in human disease. These studies are the first to highlight the molecular interplay of AR coactivators, CBP/p300, and HR-mediated repair in CRPC with preclinical and clinically relevant models (Fig. 7G). In sum, these studies nominate potential combinatorial targets for therapeutic intervention to enhance patient outcome.

Previous studies using reporter assays nominated CBP/p300 as coregulators in support of AR and c-Myc^32,33,36–40^, and in PCa models, CBP/p300 expression is induced in response to androgen ablation, suggesting that CBP/p300 may support disease progression by amplifying basal AR activity in the castration setting, and thereby enhance tumor progression. This concept is further supported by our previously published^35^ study and this current study. CBP and p300 are often treated as interchangeable due to their functional similarities. However, both proteins have nuanced distinctions that enable unique functions^58,77^. This study delves into elucidating these potential differences utilizing cistromic and transcriptomics analyses with CBP/p300 pharmacological and genomic attenuation (Figs. 3-4). Notably, this is the first study to identify the CBP and p300 cistromes in CRPC models (Fig. 3). Importantly, cistromic analyses in CRPC revealed distinct binding regions of CBP and p300 (Fig. 3B). CBP preferentially binds to the intergenic region (42%) while p300 binds to the intronic (33%) and promoter (26%) regions, which is shifted with CBP/p300 bromodomain inhibition via CCS1477 treatment. Moreover, while the transcriptional partners with which CBP/p300 interact with are similar as seen with the FOX family of transcription factors (Fig. 3C), the mechanism of action may be different as seen with the unique *de novo* motif analyses (Fig. 3D). Proximal to CBP binding, motifs are enriched for Ets-factors Elk4 and Sp1, which are linked to PCa oncogenic signaling^62,64,67–70^. Intriguingly, proximal to p300 binding, motifs are enriched for GFY, ZNF143, and HOXB13, which are linked to mitochondrial signaling and zinc-finger function^71–78^. These unique binding regions and motif analyses by the CBP/p300 paralogs could impact distinct downstream signaling and biological function as seen with the Hallmark pathway analyses indicating distinct functions of CBP and p300 (Fig. 3E). Future studies will investigate the potential exclusive, non-redundant functions of CBP and p300 as potential therapeutic targets.

CBP/p300 have been studied for decades for their HAT functions as a potential therapeutic target for treating cancer^49,56,84,85^. This exploration has uncovered aspects of the CBP/p300 and HAT potential mechanism of action. The HAT domain has been shown to regulate DDR by regulating reactive oxygen species (ROS) production and function^55,86–88^. While this study utilizes pharmacological targeting of the bromodomain of CBP/p300, we acknowledge that targeting of the HAT catalytic domains or the CH1/TAZ binding domains are alternative options. Notably, targeting these domains were not pursued since pharmacological inhibition of the HAT domain has been therapeutically unsuccessful thus far^49,55,56,58^. In our previous paper^35^, we demonstrated that bromodomain inhibition caused DNA repair de-enrichment and G2M checkpoint arrest. Similarly, genomic attenuation of CBP/p300 with isogenic pairs of CBP and p300 knockdown utilizing shRNA in CRPC models, revealed key DDR and cell cycle pathways decreased with CBP/p300 knockdown (Fig, 4). Thus, the continuity of pharmacological inhibition and genomic decrease suggest a potential mechanism of action that was further explored in this study. Intriguingly, the potential interplay of CBP/p300 and DDR could be a novel combinatorial therapy.

Transcriptomic pathway analysis, functional molecular studies, and biological validation herein revealed that CBP/p300 govern a discrete network of transcriptional programs of cancer relevance, including regulation of DNA repair processes, cell cycle control, and metabolic regulation. While there are studies detailing CBP/p300 in influencing DNA repair, cell cycle control, and metabolic regulation^55,86–93^, the role of CBP/p300 in transcriptional regulation of double-strand DNA repair regulation in CRPC models was unknown. Functional assessment herein demonstrated that CBP/p300 regulates HR-mediated repair *in vitro*, *in vivo*, and *ex vivo*. Briefly, CBP/p300 attenuation via CCS1477 or shRNA knockdown revealed in the presence of genotoxic insult resulting in double-strand break, HR-mediated repair was impaired and PCa growth decreased (Figs. 4-6). Importantly, AR is a key driver of PCa and known factor in DDR including regulation via PARP and DNAPK^20,51,52,80,94^. Thus, potential mechanism of action for CBP/p300 promoting HR-mediating repair could be through coactivator function with AR or via modulation of multiple oncogenic transcriptional factors. As described in the cistromic analyses (Fig. 3), binding enrichment was observed for several oncogenic transcription factors of established PCa relevance, including forkhead factors, ETS factors, and zinc finger components. Future studies will be directed at determining which of these factors regulate the underpinning mechanisms for CBP/p300 and HR-mediated repair.

Finally, this study further highlights the clinical importance of CBP/p300 alterations in disease progression. Findings herein revealed that CBP/p300 are strongly associated with poor outcome in human disease (Fig. 1). Also, CBP/p300 regulation of key HR factors has translational relevance with co-occurrence of *CBP* and *p300* expression with HR targets (*ATM*, *MRE11*, and *RAD50*) in metastatic PCa patients (Fig. 7). These observations are critical given that homologous recombination deficiency (HRD) impacts PCa progression and therapeutic efficacy^95–101^. Briefly, PARP inhibitors (PARPi) have been tested and approved for BRCA deficient tumors^102–106^. Thus, the role of CBP/p300 in influencing the response to PARP1/2 inhibitors should also be explored. Given that CBP/p300 are coactivators of AR and attenuation of these factors are critical for HR-mediated DNA repair, CBP/p300 status may provide further therapeutic insight into tumors that may respond to PARPi. Determining mechanisms to directly antagonize CBP/p300 function in the clinical setting may also be an important next avenue of investigation. Currently, ongoing clinical trials to determine the efficacy and tolerability of CCS1477 (Inobrodib) include evaluation in solid tumors (clinical trial identifier NCT03568656) and hematological malignancies^107^. Specifically, an open label phase I/IIa study of CCS1477 as monotherapy and in combination with AR targeting agents will lead to more insight for the role of HRD impacting efficacy. The clinical development of CBP/p300 inhibitors may face challenges as monotherapies in metastatic PCa. Therefore, drug combinations of utilizing DDR agents with CBP/p300 inhibition might prove to be more effective. Lastly, there are recently developments of the use of PROTAC as CBP/p300 degraders and examining their effect in solid tumors^108,109^. Importantly, future studies exploring mechanisms of resistance are crucial to enhance therapeutic efficacy regardless of the pharmacological means utilized to target CBP/p300 function.

Taken together, studies herein reveal new knowledge of the interplay between CBP/p300 and DNA repair in advanced PCa, wherein CBP/p300 expression is associated with poor outcome. Unbiased sequencing, molecular interrogation, and biological assessment of CBP/p300 function suggest CBP/p300 are key drivers of HR-mediated repair by revealing a novel mechanism of action in response to genotoxic insult. Pharmacological and genetic attenuation of CBP/p300 indicated that these factors impact HR efficacy and promote CRPC growth in preclinical and clinical models. Thus, targeting AR signaling via its coactivators in combination with DDR agents offers a new potential therapeutic strategy that merits evaluation in advanced PCa clinical trials to enhance patient outcome of this lethal disease.

## Methods

### Cell culture and reagents

22Rv1 and C4-2 cells were purchased from ATCC, authenticated by ATCC, and tested for mycoplasma upon thawing of cells. Doxycycline-inducible cell line models to knockdown expression of CBP and p300 (shCON, shCBP, and shP300) were developed as previously described^35^. All C4-2 derived cell lines were cultured and maintained in Improved Minimum Essential Medium (IMEM) (Thermo Fisher Scientific, 10024CV) supplemented with 5% FBS (fetal bovine serum, heat inactivated), 1% L-glutamine (2mmol/l), and 1% penicillin-streptomycin (100 units/ml). All 22Rv1 derived cell lines were cultured and maintained in Roswell Park Memorial Institute (Gibco RPMI) (ATCC, CATALOG # A1049101) supplemented with 10% FBS, 1% L-glutamine, and 1% penicillin-streptomycin. All cells were cultured at 37°C at 5% CO_2_. CBP/p300 bromodomain inhibitor was developed from CellCentric and deta^35^1l. Briefly, CCS1477 was synthesized according to processes described in the International Patent Application, publication number WO2018073586.

### Transcriptome data analysis of two mCRPC cohorts (SU2C and RMH – Royal Marsden Hospital)

Patient cohort was utilized from previously published study^35^. All patients had given written informed consent and were enrolled in institutional protocols approved by the Royal Marsden (London, UK) ethics review committee (reference no. 04/Q0801/60). Human biological samples were sourced ethically, and their research use was in accordance with the terms of the informed consent provided as detailed previously^35^. A total of 159 mCRPC transcriptomes generated by the SU2C–PCF Prostate Cancer Dream Team^110^ and 95 mCRPC transcriptomes from patients treated at Royal Marsden Hospital and Institute of Cancer Research (RMH/ICR) were analyzed^111^ using TopHat2-Cufflinks pipeline. Gene expression as fragments per kilobase of transcript per million mapped reads (FPKM) was calculated.

### Chromatin Immunoprecipitation (ChIP)-Sequencing

22Rv1 cells were plated in hormone-proficient media. ChIP was performed as previously described^15^. Briefly, cells were cross-linked with 1% fresh formaldehyde for 10 mins at room temperature. Chromatin was sheared to 200-700 bp using Active Motif Ultrasonicator for 30 cycles (30 seconds on, 30 seconds off). CBP and p300 antibodies were obtained from Cell Signaling (CST). ChIP-Seq was performed by Active Motif according to their guidelines and protocols including antibody testing, library preparation, and sequencing. ChIP-Seq data will be deposited in the NCBI GEO with the accession code upon publication.

### ChIP-Sequencing Analysis

Analyses was performed by expert bioinformaticians. Briefly, FASTQ files were assessed for quality using FASTQC v0.11.5. Reads were aligned to the human genome reference version hg19 using bowtie2 v2.3.2^71^ with default parameters. Peak calling was performed using MACS2 v2.1.1^72^ with combined replicates, utilizing a q < 0.05 cutoff. ChIP-Seq binding heatmaps and profiles were generated using deepTools v2.5.7^73^. Peak annotation and motif analysis performed using Homer v4.10.3^74^ using the parameters indicated.

### RNA-Sequencing

22Rv1 shCON, 22Rv1 shCBP, and 22Rv1 shP300 cells were treated with doxycycline for 5 days to knockdown expression of CBP or P300 in biological quadruplicate. RNA was extracted and purified using the miRNeasy kit (QIAGEN, 217004) by following manufacturer’s instructions. Novogene performed RNA-Sequencing following their company polices. Briefly, TruSeq Stranded Total RNA Library Prep Gold kit was used to construct the RNA-Seq libraries. NextSeq 500 sequencer from Illumina was utilized to sequence samples using single-end 75bp reads.

### RNA-Sequencing Analysis

FASTQ files were aligned using STAR v2.5.2a against the human genome (hg19)^112,113^. Read counts for each gene were generated using featureCounts utilizing Ensembl as reference gene annotation set^114^. Differential gene expression data were generated using DESeq2 v1.12.4^115^. Gene set enrichment analysis (GSEA) was performed using HALLMARKS and KEGG gene signatures from the Molecular Signature Database^116^. RNA-Seq data will be deposited in the NCBI GEO with the accession code upon publication.

### Flow Cytometry

All 22Rv1 and C4-2 parental and derived cells were plated at equal densities in hormone-proficient media (i.e., FBS condition). Then follow all treatment conditions described, cells were incubated with BrdU (1:1000) for 2 hours prior to harvesting. Cells were fixed and processed as previously described^117^. At least 10,000 events per sample were assessed. Analysis was performed using InCyte software (Guava) for cell-cycle profile with BrdU incorporation and PI (propidium iodide).

### Gene Expression Analysis

All 22Rv1 and C4-2 parental and derived cells were plated at equal densities in hormone-proficient media (i.e., FBS condition). Trizol (Invitrogen) was used to isolate RNA and SuperScript VILO (Invitrogen) was used to generate cDNA following manufacturer’s instructions. PowerSybr (Fischer Scientific 43-676-59) and the ABI StepOne Real-Time PCR system were utilized in accordance with manufacturer’s specifications to perform quantitative PCR (qPCR) analyses. The primers used are depicted in the table below.

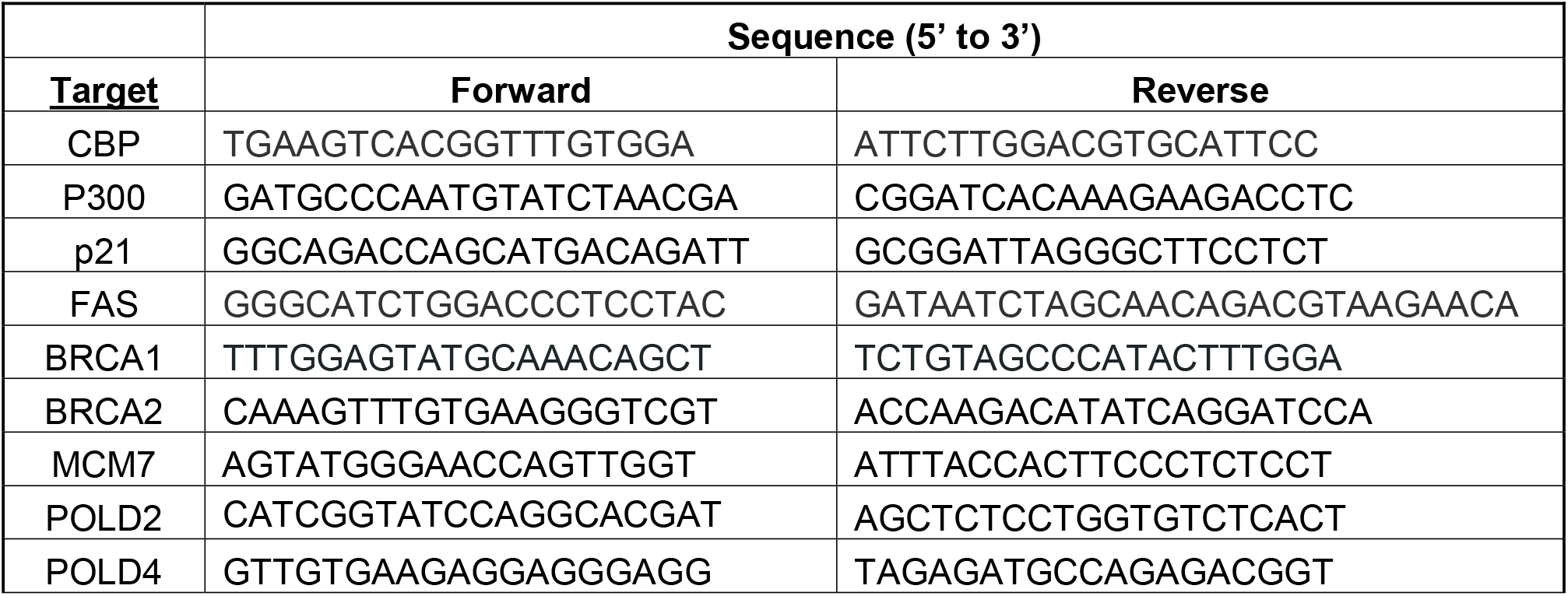
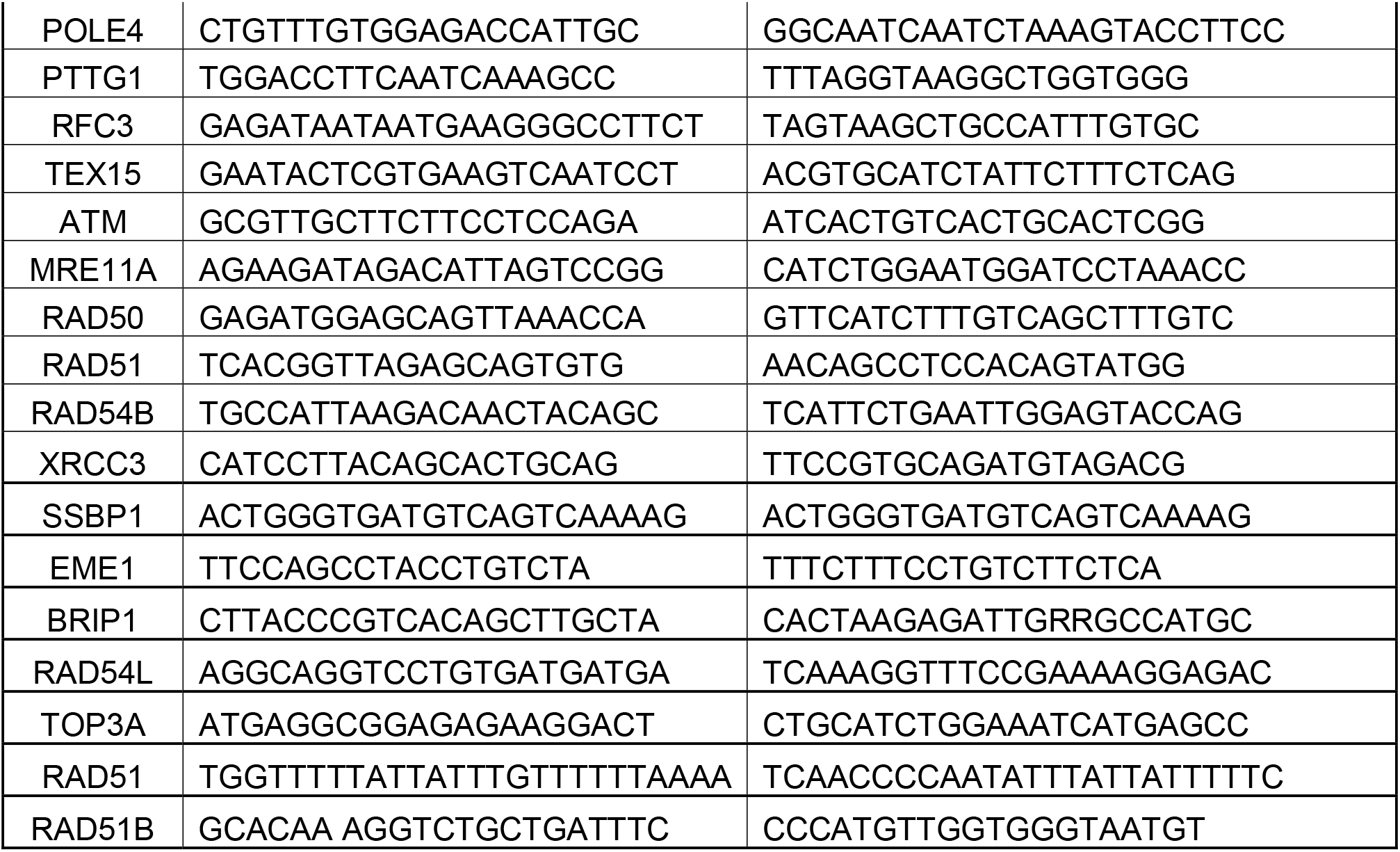

### Immunoblotting

All 22Rv1 and C4-2 parental and derived cells were plated at equal densities in (i.e., FBS condition). Generation of cell lysates was described previously^52^. 40-50 μg of lysate was resolved by SDS-PAGE, transferred to PVDF (polyvinylidene fluoride) membrane, and analyzed using the following antibodies at 1: 500 dilution – CBP (D9B6), p300 (D2X6N), 1:1000 dilution – ATM (Cell Signaling Technology 2873), phospho-ATM (Ser1981) (CST 5883S), CHK2 (Bethyl A300-619A), phospho-CHK2 (Thr68) (CST 2661T), MRE11 (CST 8344T), RAD50 (CST 8344T), RAD51 (Abcam ab63801), XRCC3 (Novus NB100-165), and 1: 5000 – Vinculin (Sigma-Aldrich V9264).

### Proliferation Assays

All 22Rv1 and C4-2 parental and derived cells were plated at equal densities in (i.e., FBS condition). Cells were treated with either IR (irradiation), CBP/p300 inhibitor (CCS1477, CellCentric)^35^, or doxycycline, which was refreshed every 48 hours. Cell number was quantified using the Quanti-IT Pico Green dsDNA assay kit (Thermo Fisher) at the indicated times of treatment.

### Xenograft Analysis

All animals were housed in pathogen-free facilities. All mouse work was approved by the Axis BioServices Animal Welfare and Ethical Review Committee and conducted under license and within the guidelines of the Home Office Animals (Scientific Procedures) Act 1986. Prostate xenograft tumors were established by subcutaneous injection of 22Rv1 cells into non-castrated male athymic nude mice. Once tumors had reached approximately 150 mm^3^, animals were randomized into control and treated groups. In all studies, tumor volume (measured by caliper), animal body weight, and condition were monitored at least twice weekly. 22Rv1 xenograft RNA was after 28 days of treatment of 20 mg/kg of CCS1477 (26 for the vehicle) and that the CP50cs were short term treated for 8 days and long term was until they reached 300% of their original growth. Tumor samples were collected for analyses of pharmacodynamics biomarkers. All the mice used in this study were male. No mice lost more than 5% of their body weight throughout the duration of the study. All mouse work was carried out in accordance with the Institute of Cancer Research (ICR) guidelines, including approval by the ICR Animal Welfare and Ethical Review Body, and with the UK Animals (Scientific Procedures) Act 1986. The CP50 PDX and CP50s PDX was derived from a metastatic lymph node biopsy from a patient with CRPC who had received all standard-of-care therapies for prostate cancer as previously described^35^.

### PDE (Patient Derived Explant)

Fresh prostate cancer specimens were obtained with written informed consent from men undergoing robotic radical prostatectomy at St Andrew’s Hospital, Adelaide, through the Australian Prostate Cancer BioResource. Dissected tissue fragments were utilized as *ex vivo PDE* cultures as previously described^53,83,118,119^. University of Adelaide’s Institutional Review Board has reviewed this protocol and deemed this research to follow federal regulations (Approval # HREC-2012-016). PDE cultures were treated with media containing CCS1477 (1 or 5 μM) or vehicle alone (DMSO) and harvested after 48 hours for immunohistochemistry (IHC) analyses of the proliferative marker, Ki67, and qPCR for gene expression analyses of HR target genes. For histological analysis, explants were formalin-fixed and paraffin embedded. For RNA analysis, explants were stabilized in RNAlater® (Ambion, TX, USA) at 4°C overnight and then stored at −80°C.

### Immunohistochemistry (IHC)

For PDEs, immunostaining of formalin-fixed paraffin-embedded sections (2-4 μm) was performed with the Bond RX automated stainer (Leica Biosystems, Germany). Antigen retrieval was 20 mins at 100°C using the Bond Epitope Retrieval solution 2 (EDTA based buffer, pH 9). FFPE sections were stained with Ki67 primary antibody (Clone MIB-1, DAKO, Denmark) (1:200) and a goat anti-mouse IgG biotinylated secondary antibody (DAKO, Denmark). using standard techniques previously described^120^.

### Homologous Recombination (HR) Activity Assay

U2OS-DR-GFP cells are a modified osteosarcoma cell line that were generated by Dr. Jasin and were utilized to assess HR activity as previously described^81,82,121^. Dr. Roger A. Greenberg (University of Pennsylvania) provided these cells that were utilized in this study. These cell lines were transfected with siCBP or siP300 using Dharmafect 4 reagent following manufacturer’s instructions. 48 hours post transfection, ISce1 plasmid was transfected into cells to induce DNA breaks. Cells were treated with ATM inhibitor (KU-55933, Sigma-Aldrich SML1109) or CBP/p300 inhibitor (CCS1477, CellCentric) for the last 16 hours of the assay. Cells were harvested and GFP positive cells were quantified via flow cytometry.

### Statistics

All experiments were performed in technical triplicate with at least 3 biological replicates per condition. Data are displayed as mean ± standard error of the mean (SEM). Statistical significance (p < 0.05) was determined using Student’s t-test, one-way ANOVA, and two-way ANOVA on GraphPad Prism Software as appropriate and indicated in applicable figure legends. For the analysis of patient biopsies, nuclear CBP and p300 protein levels were reported as median values with IQRs. For paired and/or same PCa patient expression studies, the Wilcoxon matched pair signed rank test was used to compare differences in protein expression levels. The correlation between nuclear CBP and p300 protein expression was determined using Spearman correlation. Overall survival was defined as time from diagnosis (defined above) to date of death or last follow-up/contact. Patient outcomes were compared by nuclear CBP and p300 protein expression (H-score) at diagnosis; median overall survival and median time to CRPC were estimated using the Kaplan–Meier method, and respective hazard ratios were obtained by Cox regression.

### Study approval

The use of patient and clinical material was approved by the ethical committees from each of the following institutes: the Sidney Kimmel Cancer Center at Thomas Jefferson University (Pennsylvania, USA), Institute of Cancer Research (ICR – London, United Kingdom), and the University of Adelaide (Adelaide, Australia).

## Data Availability

The datasets generated during the current study have been deposited in public repositories. RNA-Seq data will be deposited in the NCBI GEO with the accession code upon publication.

## Acknowledgements

This work was supported by a Young Investigator Award and Challenge Award from the Prostate Cancer Foundation (to AAS, AS, & KEK), NCI R01-CA182569 (KEK), the Sidney Kimmel Cancer Center (SKCC) Support Grant (5P30CA056036), CPDR-CORE Funds (AAS, XAS), Wellcome Trust Clinical Research Career Development Fellowship (AS) and the Cancer Genomics and Translational Research/Pathology core services at SKCC. LMB is supported by the South Australian Immunogenomics Cancer Institute at the University of Adelaide and Cancer Australia (ID#1138766). Additionally, we would like to thank Dr. R. Greenberg (University of Pennsylvania) for providing the U2OS-DR-GFP cells. The authors thank the study participants, urologists, nurses, histopathologists and coordinators of the Australian Prostate Cancer Bioresource that provided specimens for this research. The collaborators played a role in the design of the study; data collection; analysis; interpretation; and review and approval of the manuscript. Lastly, we would like to thank former Knudsen lab members and current CPDR colleagues for providing suggestions and support for this study.

## Author Contributions

Conceptualization, C.M.M., A.S., X.A.S, L.B., J.B., K.E.K, and A.A.S.; Methodology, C.M.M., K.E.K, and A.A.S.; Software, C.M.M, W.Y., and D.B.; Validation, S.S, C.C.M., L.R., and A.A.S.; Formal Analysis, S.S., C.M.M, S.N.C., W.Y, D.B., J.W., E.G.D., T.J., L.B., and A.A.S.; Investigation, A.A.S., M.A., T.M.S., Y.Z., A.B., N.K.R, and M.J.S.; Resources, C.M.M., A.S., M.J.S., L.B., J.B., K.F., N.B., N.P., K.E.K, and A.A.S.; Writing, S.S. and A.A.S; Writing – Review & Editing, S.S., C.M.M., A.S., M.J.S., L.B., J.B., K.F., N.B., N.P., K.E.K, and A.A.S; Visualization, S.S., C.M.M., W.Y., D.B., L.B., J.D., A.A.S.; Funding Acquisition, K.E.K, and A.A.S.

## Competing Interests Statement

The following are disclosures for authors on this manuscript: Karen E. Knudsen is the CEO of American Cancer Society (ACS). Adam Sharp has received travel support from Sanofi, Roche-Genentech and Nurix, and speaker honoraria from Astellas Pharma and Merck Sharp & Dohme. He has served as an advisor to DE Shaw Research & CHARM Therapeutics and has been the CI/PI of industry-sponsored clinical trials. No disclosures for the other authors. Neil Pegg, Nigel Brooks, and Kris Frese are employees and shareholders of CellCentric Ltd. Neil Pegg is also a board director and the inventor on CCS1477 patents. Johann S de Bono reports grants from CellCentric during the conduct of the study; grants and personal fees from Daiichi Sankyo, AstraZeneca, Pfizer, Bayer Oncology, MSD, Merck Serono, Harpoon, and Genentech/Roche, personal fees from Eisai and Constellation, and grants from Sun Pharma outside the submitted work; in addition, Johann S. de Bono has a patent for Abiraterone licensed and with royalties paid from Janssen. No other disclosures were reported.

