## Supplemental Figures for "AR coactivators, CBP/p300, are critical mediators of DNA repair in prostate cancer"

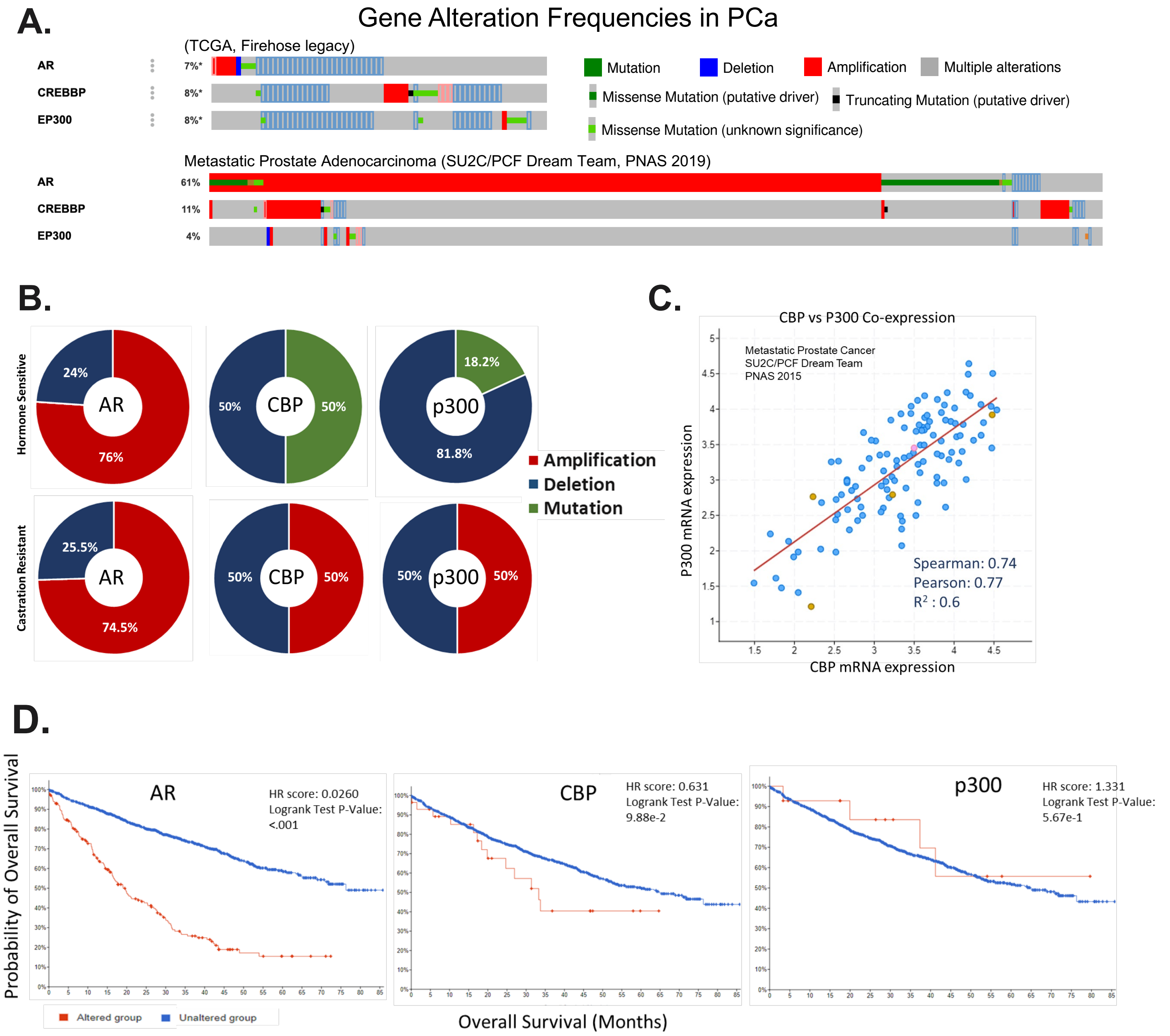


**Supplementary Figure 1. Altered CBP/p300 expression correlates with aggressive disease. A.** Frequency of AR, CBP, and p300 alterations in primary PCa. **B-C.** Correlation of CBP and p300 mRNA expression in metastatic PCa patient tumors from the Prostate Adenocarcinoma (MSK/DFCI, Nature Genetics 2018 and Metastatic Prostate Cancer (SU2C/PCF Dream Team, Cell 2015) data sets from cBioportal. Hormone sensitive PCa total alterations AR n= 167, CBP n=26, and p300 n=11, and castration-resistant PCa AR n=94, CBP n=8, and p300 n=2. **D.** cBioportal analyses for overall survival from the Prostate Adenocarcinoma (MSK, Clin Cancer Res. 2022) data set.

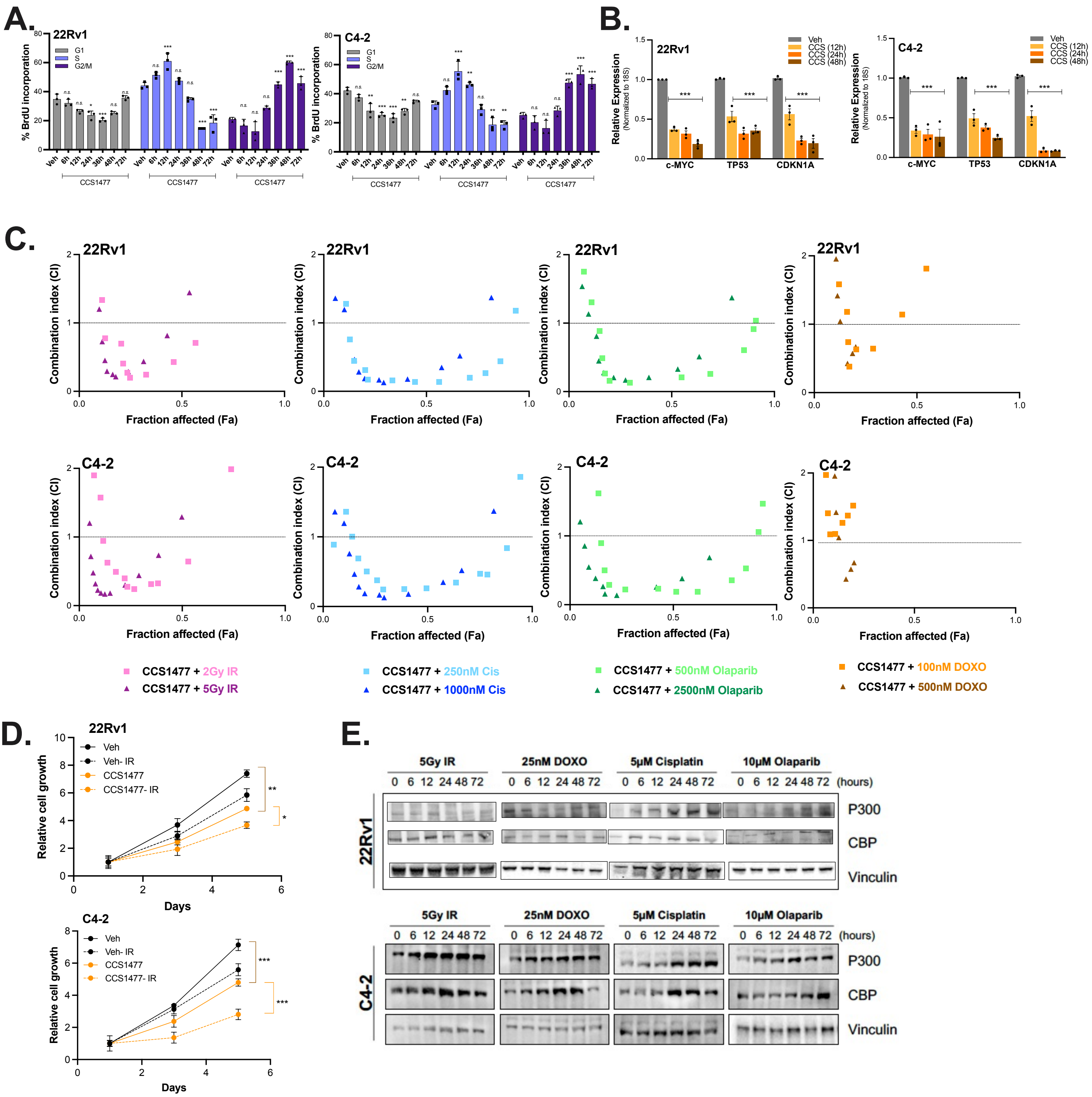

**Supplementary Figure 2. CCS1477 sensitizes CRPC cells to AR-dependent DNA repair. A-B.** 22Rv1 and C4-2 cells treated with CBP/p300 inhibitor (CCS1477) for increasing time (4-72 hours). **A.** Cells were harvested and analyzed for cell cycle analyses with flow cytometry. **B.** Cells were harvested, RNA isolated, and qPCR analyses of key target genes. **B.** Cells were harvested and analyzed for cell cycle analyses with flow cytometry **C.** 22Rv1 (top) and C4-2 (bottom) cells were treated with increasing dosages of indicated DDR agents (IR, cisplatin, olaparib, or doxorubicin) with or without CCS1477, and drug sensitivity assays were performed on Day 5 using Picogreen. Combination Index (CI) analyses were performed for conditions performed above. **D.** 22Rv1 (top) and C4-2 (bottom) cells were treated with or without CCS1477 (250 nM) and IR (5 Gy) at Day 0. Cells were harvested for growth assays analyses on Days 0, 3, and 5 using Picogreen. **E.** WB analyses of AR, CBP, p300, c-MYC, and Vinculin in response to IR, cisplatin, olaparib, and doxorubicin in 22Rv1 and C4-2 cells. n=3, *p<0.05, **p<0.01, & ***p<0.001.

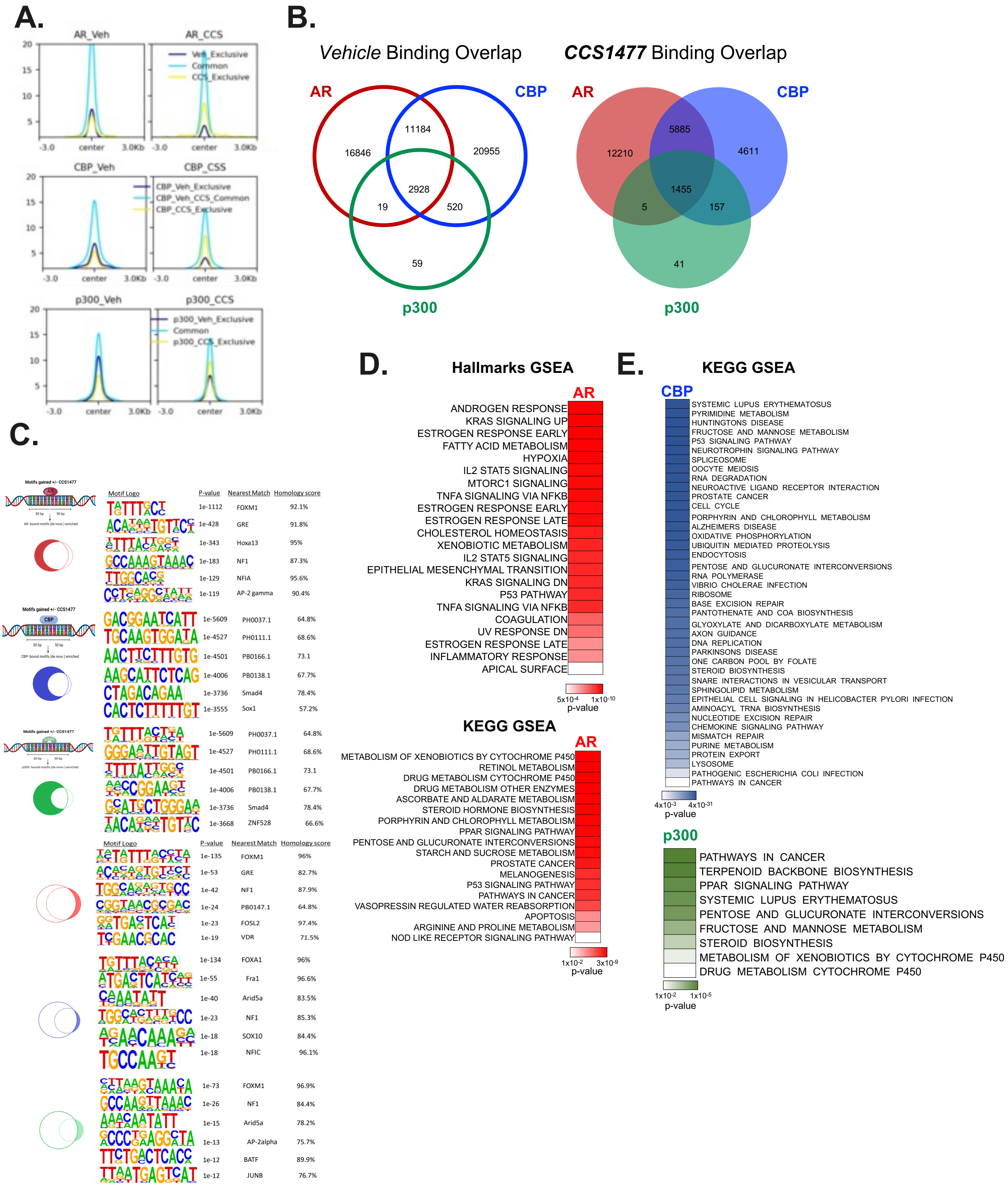


**Supplementary Figure 3. CBP/p300 inhibition drastically reduces the AR , CBP, p300 cistromic landscape. A.** Overview of bindings sites of AR, CBP, & p300 in Vehicle treated 22Rv1 cells. **B.** Overlap of bindings sites of AR, CBP, & p300 in Vehicle condition and CCS1477 treated condition in 22Rv1 cells. **C.** Motifs of binding regions of AR, CBP, and p300 exclusive to Veh or CCS1477 conditions visualized for top 6 highest p-values. **E.** Hallmarks and KEGG GSEA pathways regulated by AR. **E.** KEGG GSEA pathways regulated by CBP and p300 in CCS1477 treated 22Rv1 cells.

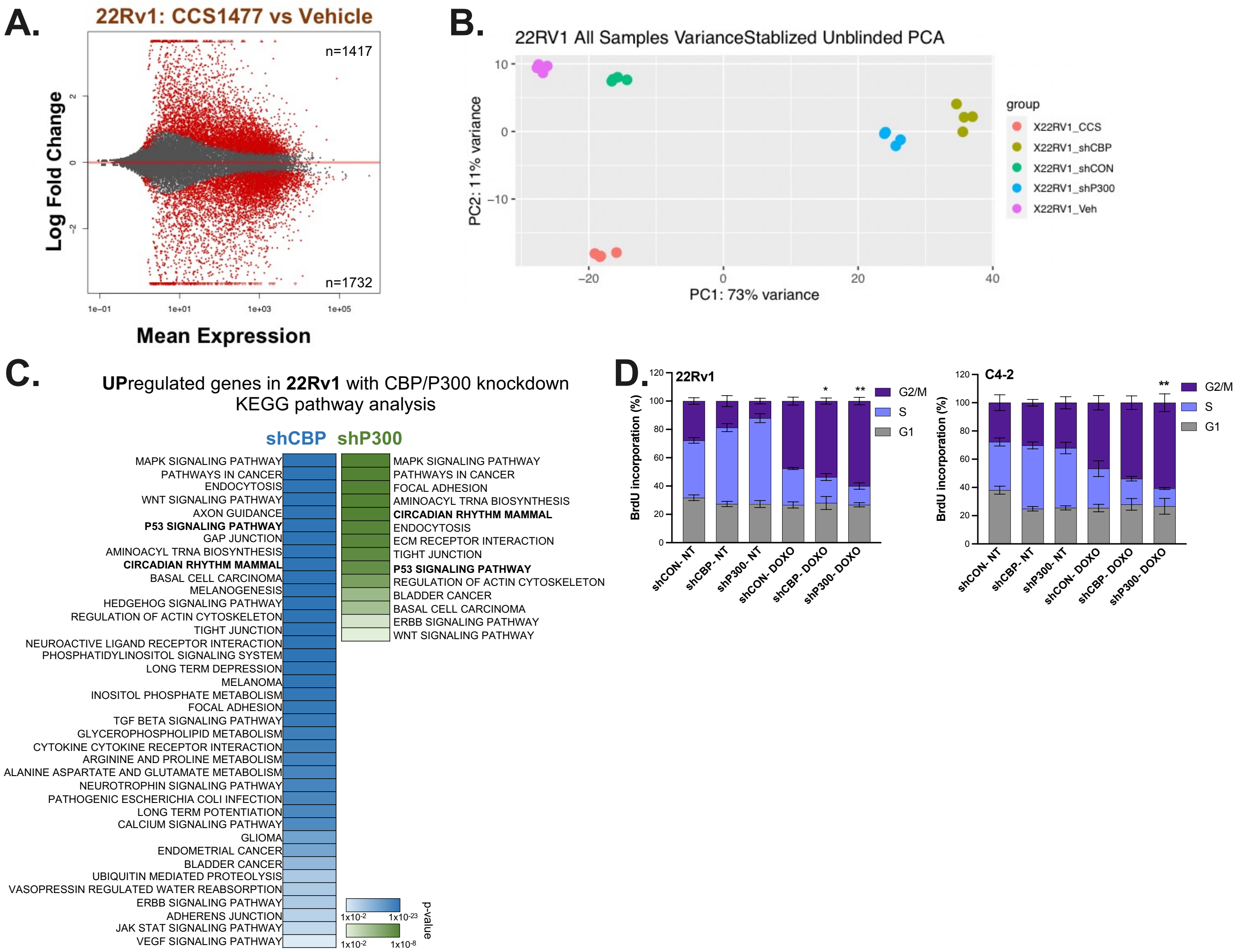


**Supplementary Figure 4. CBP/p300 bromodomain knockdown alters global transcriptome and regulates cell cycle & DDR. A.** 22Rv1 cells treated with Vehicle or 100 nM CCS1477 treatment for 24 hours as detailed in previously published data in Welti *et al*, Cancer Discovery 2021. MA plot depicts gene expression modulation with the number of significant upregulated and downregulated transcripts with p<0.001 and fold change (FC)> 2.0. **B.** PCA blot of RNA-Seq analysis was performed on quadruplet samples for 22Rv1-shCON, 22Rv1-shCBP, and 22Rv1-shP300 in addition to 22Rv1 with CCS1477 treatment. **C.** GSEA pathway analyses (KEGG) of upregulated in 22Rv1 cells with CBP and p300 knockdown treatment using FDR<0.25, p<0.0001, FC>2. **D.** C4-2 and 22Rv1-shCON, -shCBP, and –shp300 cells were treated with doxycycline to knockdown CBP and p300. Then cells were treated with 10 nM Doxorubicin (DOXO). Flow analysis was performed to determine changes in cell cycle progression.. n=3, *p<0.05, **p<0.01, & ***p<0.001.

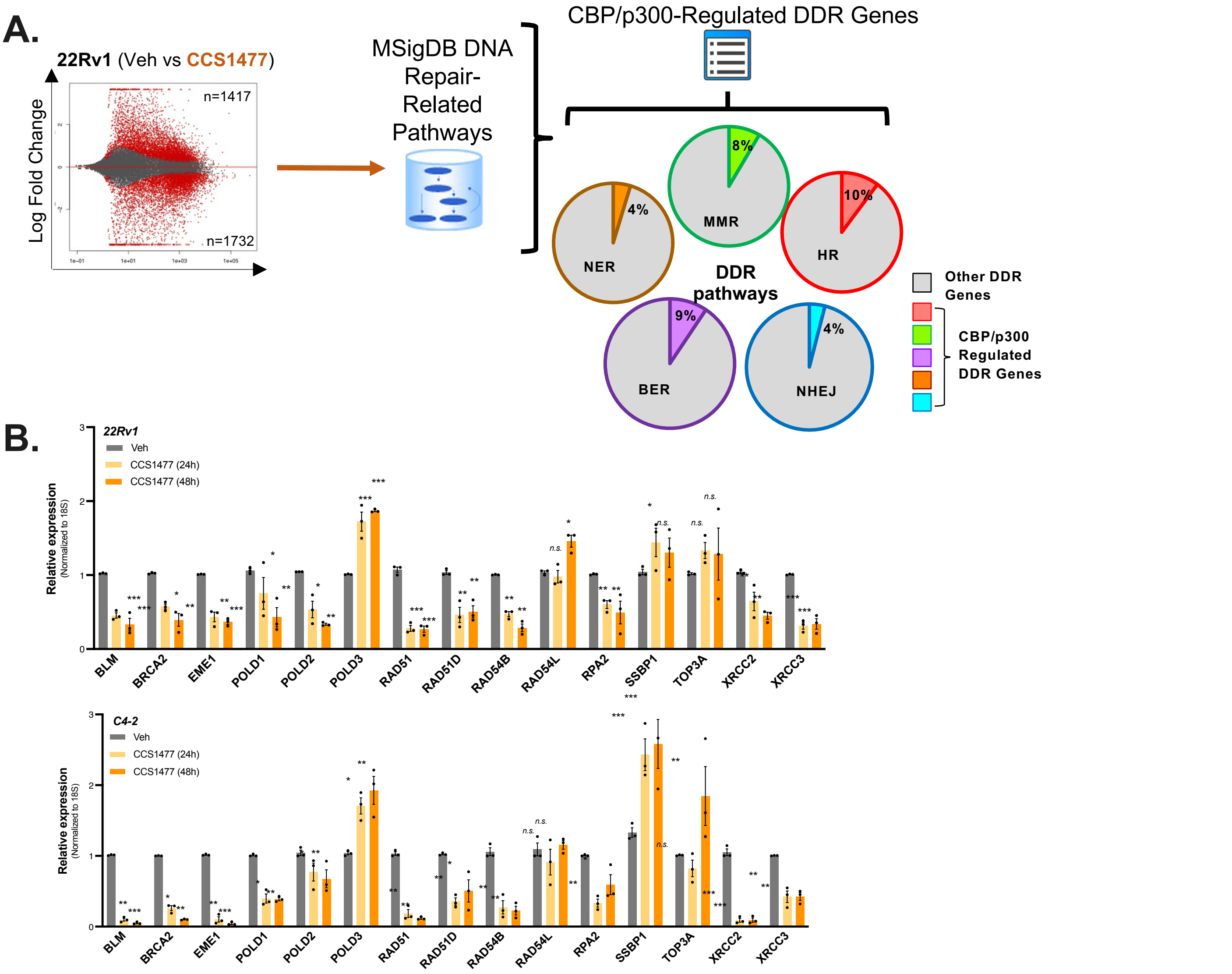


**Supp Figure 5. CBP/p300 inhibition impacts double-strand DNA break repair. .** Schematic describing the comparison of RNA-Seq data sets. Briefly, transcriptions with p<0.001 and fold change (FC) > 2.0 were identified and then organized into specific DDR pathways. **B.** Validation of HR gene targets in CRPC cells with increasing time of CCS1477 treatment. n=3, *p<0.05, **p<0.01, & ***p<0.001.

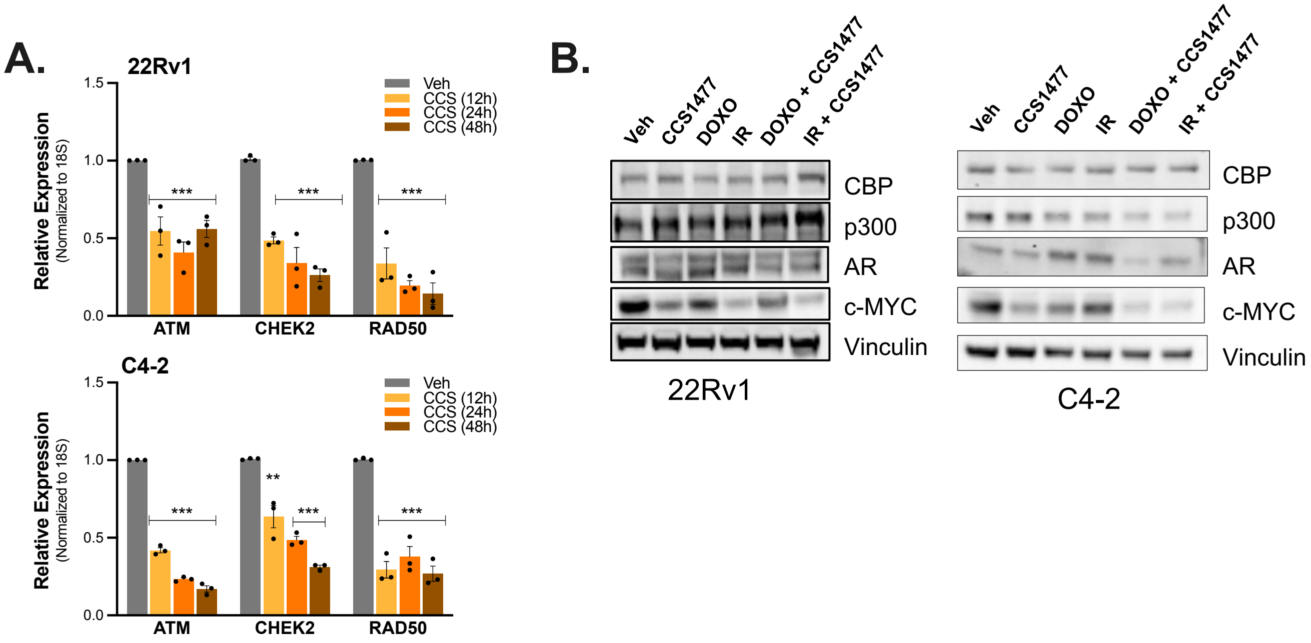


**Supp Figure 6. CBP/p300 inhibition and knockdown decreases HR efficiency. A.** Validation of HR gene targets in CRPC cells with increasing time of CCS1477. **B.** 22Rv1 & C4-2 cells were treated with with CBP/p300 inhibitor with or without genotoxic insult of 5 Gy IR or 10 nM Doxorubicin treatment for 24 hours. Changes in HR factors protein expression were analyzed. n=3, *p<0.05, **p<0.01, & ***p<0.001.

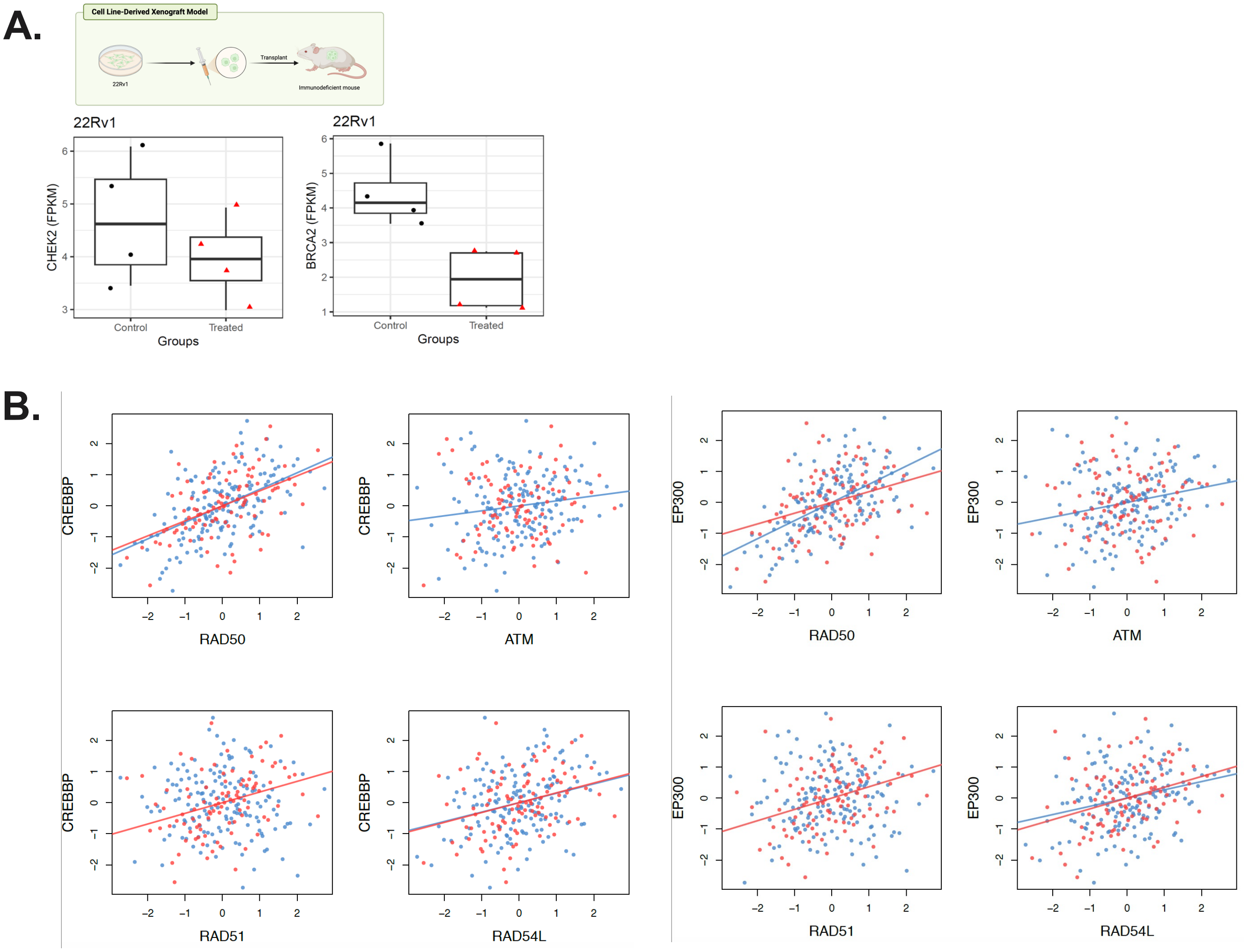


**Supp Figure 7. CBP/p300 inhibition mediates HR repair and impacts patient outcome. A.** HR expression in 22Rv1 xenografts with CCS1477 (20mg/mL). Tissue harvested, RNA isolated, and analyzed for HR gene expression. **B.** Correlation of HR genes (RAD50, ATM, RAD51, RAD54L) with CBP and EP300 mRNA expression in patient cohorts.
